# Taking full advantage of modelling to better assess environmental risk due to xenobiotics

**DOI:** 10.1101/2021.03.24.436474

**Authors:** Sandrine Charles, Aude Ratier, Virgile Baudrot, Gauthier Multari, Aurélie Siberchicot, Dan Wu, Christelle Lopes

**Affiliations:** Université de Lyon, Université Lyon 1, CNRS UMR5558, Laboratoire de Biométrie et Biologie Evolutive, 69100 Villeurbanne, France

**Author notes:** This work was performed using the computing facilities of the CC LBBE/PRABI. This work benefited from the French GDR “Aquatic Ecotoxicology” framework which aims at fostering stimulating scientific discussions and collaborations for more integrative approaches. This work is part of the ANR project APPROve (ANR-18-CE34-0013) for an integrated approach to propose proteomics for biomonitoring: accumulation, fate and multi-markers (https://anr.fr/Projet-ANR-18-CE34-0013). These two authors equally contributed.

**Keywords:** dose-response models, bioaccumulation factors, toxicokinetic-toxicodynamic model, uncertainty, accessibility

## Abstract

In the European Union, more than 100,000 man-made chemical substances are awaiting an environmental risk assessment (ERA). Simultaneously, ERA of chemicals has now entered a new era. Indeed, recent recommendations from regulatory bodies underline a crucial need for the use of mechanistic effect models, allowing assessments that are not only ecologically relevant, but also more integrative, consistent and efficient. At the individual level, toxicokinetic-toxicodynamic (TKTD) models are particularly encouraged for the regulatory assessment of pesticide-related risks on aquatic organisms. In this paper, we first propose a brief review of classical dose-response models to put into light the on-line MOSAIC tool offering all necessary services in a turnkey web platform whatever the type of data to analyze. Then, we focus on the necessity to account for the time-dimension of the exposure by illustrating how MOSAIC can support a robust calculation of bioaccumulation factors. At last, we show how MOSAIC can be of valuable help to fully complete the EFSA workflow regarding the use of TKTD models, especially with GUTS models, providing a user-friendly interface for calibrating, validating and predicting survival over time under any time-variable exposure scenario of interest. Our conclusion proposes a few lines of thought for an even easier use of modelling in ERA.

**Graphical art:** 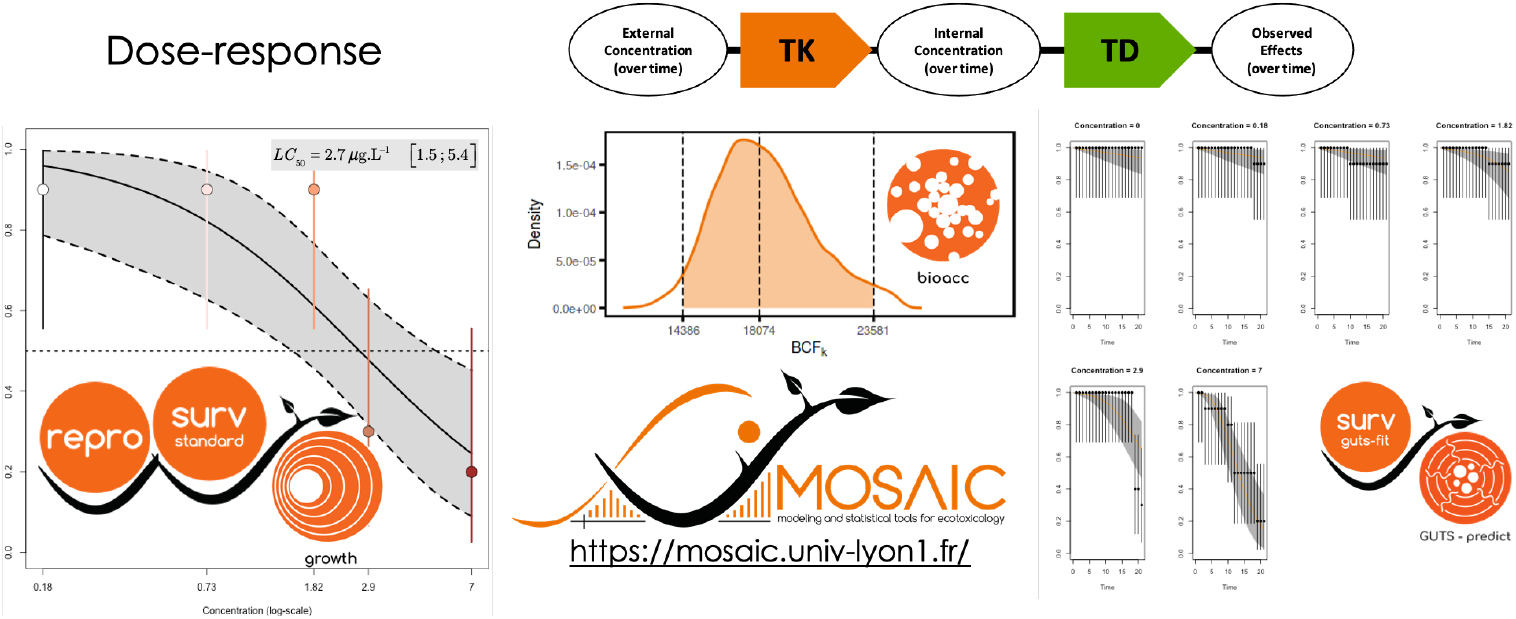

## 1 Introduction

Effects of contaminants may occur at all levels of biological organization, from molecular to ecosystem-level responses (Clements, 2000). From one level to the next the answers to exposure may strongly differ, from DNA damage metabolism disorders to loss of biodiversity or changes in food web structures. Hence, an effective translation of information through increasing organization levels (e.g., from individual to population) will provide more ecologically relevant endpoints as stated by the adverse outcome pathway concept (Ankley et al., 2010), together with increased temporal and spatial scales of the underlying processes. At the opposite, going down at inferior levels of biological organisation is crucial to finely decipher the underlying mechanisms and their specificity (Preuss et al., 2009). From the molecular to the ecosystem scales, each individual, population and community levels may appear to be the best compromise between ecological relevance and understanding of mechanisms. This explains why the vast majority of mathematical models focus on a specific biological scale, while few allow for extrapolation between these levels.

Whatever the level of biological organization, there are challenges for which mathematical models are or will be crucial. At the community level, we can mainly distinguish two categories of models. Some models consider a community as a set of species chosen to be representative of a given ecosystem without modelling the between-species interactions; this is the case with species sensitivity distributions (SSD), based on fitting probability distributions. They are used in ERA for extrapolating among species and across levels of bio-logical organization, but they are overly simplistic and likely to lead to both over-estimates and under-estimates of risk (Forbes & Calow, 2002; Forbes & Galic, 2016). Other models, based on ordinary (ODE) or partial (PDE) differential equations, will aim to describe the community functioning accounting for all types of ecological interactions as done for example by AQUATOX, the simulation model for aquatic systems from US EPA (Park et al., 2008).

At the population level, the key issue is to include individual effect models to refine the prediction of population dynamics. Indeed, effects of chemical substances do not depend only on exposure and toxicity, but also on factors such as life history characteristics and population structure. Population models are also helpful to identify critical demographic traits regarding given species-compound combinations. As reviewed in Schmolke et al. (2010), population models are mainly based on ODE/PDE, projection matrices or individual-based approaches. Although a broad range of these ecological models is available in the scientific literature, they are still rarely used in support of regulatory ERA (Schmolke et al., 2010), probably due to their inherent complexity and a lack of easy tools in order to run them, except home-made computer codes rather designed for specialists.

In this paper, we focus on the individual level, where modelling has been prominent for a long time already with dose-response (DR) models providing toxicity values (namely, standard lethal *LC*_*x*_ or effective *EC*_*x*_ concentrations) allowing to identify critical life history traits for given species-compound combinations (Ritz, 2010). Nevertheless, scientific knowledge still remain poor re-garding the physiological modes of action of compounds and how they vary across species and compounds (Ashauer & Jager, 2018). Additionally, authorities today recognize the need to account for the time-dependency of the effects to better assess risk under complex exposure situations (e.g., environmentally realistic concentrations, various exposure routes, biotransformation processes, mixture effects). To this end, the toxicokinetics (TK) and the toxicodynamics (TD) of the effects require to be modelled. The TK part relates the exposure concentration to the internal concentration within organisms, considering various processes such as accumulation, depuration, metabolization and excretion (ADME). TK models are typically used to calculate bioaccumulation factors from data collected in standard bioaccumulation tests (OECD, 2012) and new perspectives are offered by a recent modelling approach (Ratier et al., 2019) associated with a ready-to-use tool (Ratier et al., 2020). The TD part makes the link between damages suffered by organisms due to internal bioaccumulated concentrations with observable effects on life history traits such as an increased mortality or a reduced growth. Combined TKTD models are recommended by EFSA to refine Tier-2 risk assessment, especially for plant protection products acting on aquatic organisms when exposed to time-variable exposure profiles (European Commission, 2013; Ockleford et al., 2018; Brock et al., 2020). In particular, the EFSA already considers ready-to-use for ERA the TKTD models dedicated to the prediction of survival over time, and the EFSA encourages more research for the other types of TKTD models, namely those based on the Dynamic Energy Budget (DEB) theory for growth and reproduction of ectotherm species and those for macrophytes. The reason why General Unified Threshold models for Survival (GUTS models) are already operable in support of the daily work of regulators is the availability of a general framework that unify all of survival models, as well as easily accessible, user-friendly and transparent turnkey tools, allowing to run them with only several user actions. Tools for GUTS models are also known to provide reproducible results, without the need for the users to invest in underlying mathematical and statistical aspects (Jager & Ashauer, 2018).

Among available modelling tools dedicated to ecotoxicity, the MOSAIC platform proposes a suite of services within an all-in-one web site. MOSAIC is an acronym for MOdelling and StAtistical tools for ecotoxICology, that can be accessed through any Internet browser at https://mosaic.univ-lyon1.fr/ (MOSAIC, 2013). Available since 2013, MOSAIC first proposed a service for SSD analyses via MOSAIC_*SSD*_ (Kon Kam King et al., 2014; MOSAIC-ssd, 2013). In 2014, two additional services, namely MOSAIC_*surv*_ (MOSAIC-surv, 2014) and MOSAIC_*repro*_ (MOSAICrepro, 2014) (details in Charles et al. (2018)), were offered to estimate classical toxicity values from standard survival and reproduction data, respectively, providing LC_*x*_ and *EC*_*x*_. In 2018, a new facility was integrated allowing to calibrate, validate and predict survival from GUTS models under time-variable exposure profiles: MOSAIC_*GUTS−fit*_ (MOSAICguts-fit, 2018) in combination with MOSAIC_*GUTS−predict*_ (MOSAICguts-predict, 2018; Baudrot et al., 2018b). At last, in 2020, two last services were offered: (i) MOSAIC_*growth*_ (MOSAICgrowth, 2020) delivering *EC*_*x*_ estimates from standard continuous data (such as length, weight, growth rate,…), making then available a full suite of services for standard analyses whatever the type of data collected via standard toxicity tests (Charles et al., 2021); (ii) MOSAIC_*bioacc*_ (MOSAICbioacc, 2020) fitting a variety of TK models accounting for several routes of exposure, several elimination processes and several phase-I metabolites from one parent compound (Ratier et al., 2019), from which bioaccumulation factors are automatically derived (Ratier et al., 2020). All MOSAIC modules make available a collection of example data sets, allowing new users to practice using the various features.

The purpose of this article is to present all the features of MOSAIC in order to guide academics, manufacturers and regulators to benefit from advanced and sound models in ERA in support of their daily work, meeting all expectations in terms of regulatory requirements. The first section gives insights on classical DR analyses, focusing on the last new-born service MOSAIC_*growth*_. The second section illustrates how to get bioaccumulation factors from TK models, with a focus on the selection of different models to be compared, and how to fulfil the EFSA workflow regarding the use of GUTS models for ERA (Ockleford et al., 2018). The last section aims at convincing the reader of the added-value of GUTS models for Tier-1 risk assessment when *LC*_*x*_ are required. Finally, the conclusion proposes concrete lines of thought to make the use of modelling in environmental risk assessment even easier.

### 2 Classical dose-response modelling

#### 2.1 Few words about modelling

When performing standard analyses of toxicity test data in MOSAIC, the mean tendency of the relationship between the observed endpoints and the tested concentrations is first described by a 3-parameters log-logistic model written as follows:

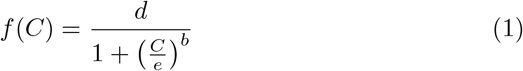

where *C* stands for the tested concentration, parameter *b* is a shape parameter translating the intensity of the effect, *d* corresponds to the endpoint value in control data (*i*.*e*., in absence of contaminant) and *e* corresponds to the *EC*_50_, that is the *C* value leading to 50% of effect compared to the control (*i*.*e*., compared to parameter *d*): 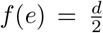. Equation (1) also assumes that lim_*C→*+*∞*_ *f* (*C*) = 0.

Then, depending on the endpoints that are observed, the variability around the mean tendency is described by an appropriately chosen probability distribution. Quantal (or binary) data (*e*.*g*., survival data) are associated with a binomial distribution. Count data (*e*.*g*., reproduction data) are associated with a Poisson distribution, possibly combined with a Gamma distribution in case of over-dispersion. Quantitative continuous data, namely data with a unit such as length or weight for example, are associated with a Normal (Gaussian) distribution. For example, in case of quantitative continuous data, the final model writes as follows:

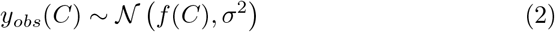

where *y*_*obs*_(*C*) stands for observations at concentration *C, f* (*C*) for the deterministic part (equation1) and *σ* for the standard deviation of the Normal law 𝒩.

Such a writing means that a total of four parameters must be estimated from observed data: *b, d, e* and *σ*. Within MOSAIC, except in MOSAIC_*SSD*_, all parameters are inferred under a Bayesian framework requiring to define prior distributions on parameters. These are automatically provided by MOSAIC based on the experimental design associated with the data as uploaded by the user. Prior distributions are then combined to the likelihood (whose writing depends on the probability law chosen to describe the variability within the data) to finally provide the joint posterior probability distribution informing on parameter estimates, their uncertainty and their correlations. Both modelling and inference processes are run automatically in MOSAIC without any action from the user to get the final results, except a single click. More information about modelling is available in Charles et al. (2018); Baudrot et al. (2018a); Charles et al. (2021); Ratier et al. (2020). MOSAIC also provides detailed information via several links: a modelling tutorial for MOSAIC_*surv*_ and MOSAIC_*repro*_ at https://cran.r-project.org/web/packages/morse/vignettes/modelling.pdf, for MOSAIC_*growth*_ at http://lbbe-shiny.univ-lyon1.fr/mosaic-growth/vignette.pdf and for MOSAIC_*bioacc*_ at http://lbbe-shiny.univ-lyon1.fr/mosaic-bioacc/data/user_guide.pdf, respectively. The subsection below illustrates how to perform to a standard DR analysis from MOSAIC_*growth*_. MOSAIC_*growth*_ has been developed in R (R Core Team, 2021) within a Shiny environment (Chang et al., 2021)

### 2.2 MOSAIC_*growth*_

Measuring growth of organisms (*e*.*g*., length of shoots, dry weight of plants, algal growth rate, size of daphnids) consists in collecting continuous quantitative data to be fitted with a DR model. MOSAIC_*growth*_ provides all useful outputs of the fitting process to check the relevance of the results, among which estimates of the effective concentration for several *x*% of interest, typically a table of *EC*_*x*_ (or *x*% Effective Rates in the field of non-target terrestrial plants). A total of 13 example data sets, concerning various species-compound combinations, are provided for new users to practice.

MOSAIC_*growth*_ makes it possible to analyse one or several data sets simultaneously (Figure 1.A), by default at the last exposure time. Regarding *EC*_*x*_ estimates, MOSAIC_*growth*_ output is the posterior probability distribution of the last *EC*_*x*_ requested by the user, as well as a summary table of all *EC*_*x*_ estimates if several of them have been requested by the user (1.B). This table includes not only the median and the 95% uncertainty interval of the *EC*_*x*_ estimates, but also censored *EC*_*x*_ values determined by taking into account the uncertainty on the estimate relatively to the range of tested concentrations (see Charles et al. (2021) for details, or http://lbbe-shiny.univ-lyon1.fr/mosaic-growth/vignette.pdf). These censored *EC*_*x*_ values can further be used for SSD analyses with MOSAIC_*SSD*_ (Kon Kam King et al., 2014).

**Fig. 1.**
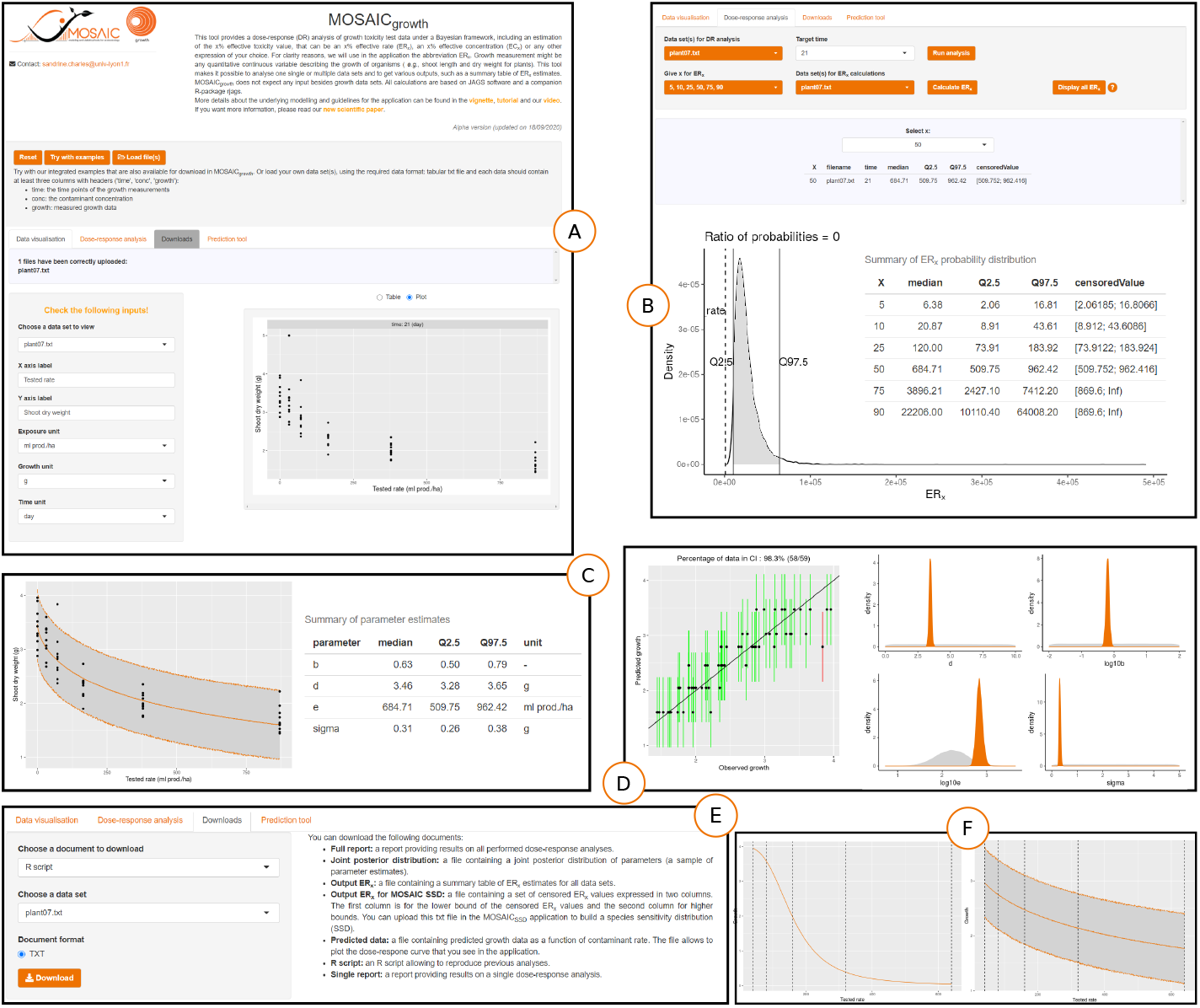
Selected pieces from the MOSAIC_*growth*_ web interface during DR analysis with the data set `plant07`: (A) upload of experimental data and visualization; (B) *ECx* estimates for *x* = 5, 10, 25, 50, 75 and 90% obtained from the results of the DR model fit and graphical representation of the probability distribution of the *EC*_90_; (C) fitted model superimposed to the observed data: median curve (solid orange line) and its uncertainty (gray area delimited by orange dotted lines) with a summary of the estimated parameters; (D) example of two model fit criteria provided by the web interface (left: ‘Posterior Predictive Check’ (PPC); right: priors and posteriors); (E) result downloading; and (F) examples with the prediction tool for a series of concentrations (40, 80, 160, 320 and 640) (left: parameters not distributed; right: distributed parameters obtained from a previous DR analysis performed with MOSAIC_*growth*_).

MOSAIC_*growth*_ also provides a visualization of the DR fit at the chosen exposure time (1.C). A table summarizes parameter estimates given as median values and their 95% uncertainty interval. In addition, goodness-of-fit criteria are provided (1.D) associated with short explanations on what is expected, in order to guide the user in checking the relevance of its results. A full tutorial is also available at http://lbbe-shiny.univ-lyon1.fr/mosaic-growth/Tutorial.pdf, especially the appendix where “no ideal” situations are presented in support of this check. In order to ensure full transparency and reproducibility of analyses, MOSAIC_*growth*_ offers the possibility of downloading various types of document, including the entire R code (1.E).

Finally, MOSAIC_*growth*_ offers a prediction tool to simulate a DR model and predict the expected relationship between a range of concentrations that the users may choose and what they can potentially achieve as effect at the final time of their experiment (1.F). Such a tool can be particularly helpful in designing future experiments for a given species-compound combination.

## 3 Accounting for the time-dependency of the effects

From a modelling point of view, the better way to account for the time-dependency of the effects is the use of TKTD models relating the exposure concentration to effects on individual life history traits via a more or less refined description of the internal damages within organisms. TKTD models allow to understand rather than to describe effects as built from underlying mechanisms. TKTD models provide time-independent toxicity parameters (as for example a no effect concentration), with outputs independent on both the experimental design and the exposure duration. TKTD models also allow to deal with time-varying exposure and to make predictions for untested situations. Above all, TKTD models allow to account for all collected data over time, while standard DR analyses only focus on a given target time (usually, the last exposure time). Section 4 will show how this may be of crucial importance for ERA.

All TKTD models can be presented according to a general scheme (Figure 2). Their specificities are related to the way both TK and TD parts are defined. Regarding TK models, all are compartment models based on ordinary differential equations, with one (the organism as a whole) or more compartments depending on their refinement. When several compartments are involved in TK models, different types are considered: either fictitious compartments (TK compartment models) or each compartment corresponding to a specific organ (physiologically-based (PB) TK models). Regarding the TD part, the type of models depends on the described endpoints: effects on survival (lethal effects) may be described by GUTS models, effects on plant growth (e.g., on macrophyte growth rate) may be described by plant models, while effects on growth and reproduction may simultaneously be described by toxicity models derived from the Dynamic Energy Budget (DEB) theory, that is DEBtox models (Ockleford et al., 2018).

**Fig. 2.**
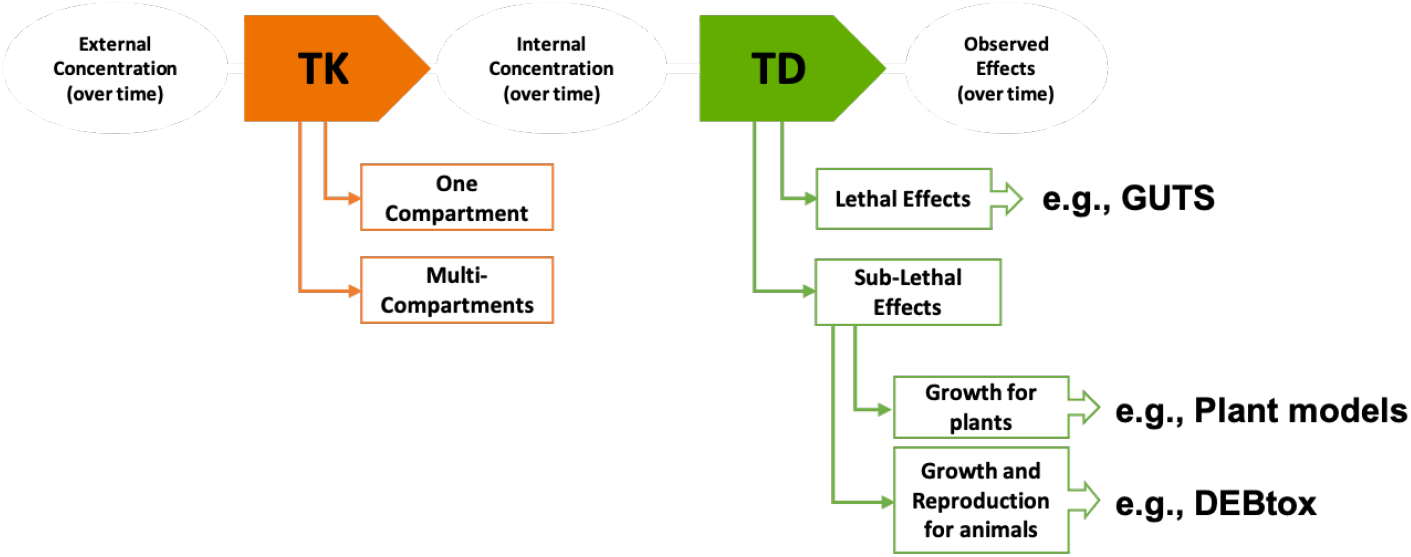
A general scheme of toxicokinetic (TK) and toxicodynamic (TD) models; GUTS stands for the General Unified Threshold model of Survival, while DEBtox stands for toxicity models derived from the Dynamic Energy Budget (DEB) theory (from Ockleford et al. (2018)).

### 3.1 TK models

#### 3.1.1 Few words about TK modelling

Chemicals are becoming potentially toxic if they bioaccumulate into the body of organisms and after being transported to a target site where they will exert effects. Chemicals may also undergo biotransformation into metabolites, which may be more or less toxic themselves. And chemicals may be eliminated from the body of organisms, for example by faeces or a phenomenon of dilution by growth. All compartment TK models assume that chemicals are evenly distributed within the compartment(s) what simplifies equations.

The most complete and complex TK models are PBTK models associating compartments to organs or physiological fluids (*e*.*g*., blood) and describing in very details all chemical fluxes between compartments; they are mostly available for aquatic species such as fish species and a number of chemical classes including plant protection products, metals, persistent organic pollu-tants, nano-particles (see Grech et al. (2017) for a review). The simplest TK model has one compartment that corresponds to one organism, in which chemicals enter (at rate *k*_*u*_) and from which chemicals are eliminated (at rate *k*_*e*_). This only-one compartment TK model will basically consider one exposure route and one elimination process. In the regulatory ERA, such models are fitted to data collected during bioaccumulation tests, which consists in an accumulation phase followed by a depuration phase. Estimates of parameters *k*_*u*_ and *k*_*e*_ are then used to calculate bioaccumulation factors (OECD, 2012).

Nevertheless, even if the most complex TK models are not always required, the very simple one reveals very limited when chemicals are present in several media, so that organisms may be exposed via several routes, and/or when several processes of elimination need to be accounted for, especially when a parent compound may biotransform into metabolites. Such situations today benefit from both a unified modelling framework (Ratier et al., 2019) and a ready-to-use tool, MOSAIC_*bioacc*_ (Ratier et al., 2020). The section below illustrates the use of the last updated version of MOSAIC_*bioacc*_.

#### 3.1.2 MOSAIC_bioacc_

MOSAIC_*bioacc*_ is a newly offered service in MOSAIC since 2020 which has been developed in R (R Core Team, 2021) within a Shiny environment (Chang et al., 2021). It allows the estimation of bioaccumulation factors associated with their uncertainty from the fit of a TK model, with only one compartment corresponding to the whole organism but several exposure routes and several elimination processes may be accounted for^1^. The model is automatically built according to the accumulation-depuration data uploaded by the user (Figure 3.A). By a single click, the user first obtains the posterior probability distribution of the kinetic bioaccumulation factor (Figure 3.B), summarized with its median and its 95% uncertainty interval (bounded by the 2.5% and 97.5% percentiles of the posterior distribution, Figure 3.C). The uploaded data may come from different types of experiments in which different routes of exposure are considered (*e*.*g*., surface water, pore water, sediment, food), as well as different elimination processes (*e*.*g*., excretion, biotransformation and growth dilution). Fitting results are plotted (Figure 3.D) superimposed to the data for the parent and its metabolites (if concerned). TK model parameters (*e*.*g*., *k*_*u*_ and *k*_*e*_ in the most simple situation) are also provided as medians and 95% uncertainty intervals (Figure 3.E). Then automatically come a number of goodness-of-fit criteria to guide the users in checking the relevance of their results (Figure 3.F). MOSAIC_*bioacc*_ provides the same goodness-of-fit criteria as MOSAIC_*growth*_, also with a short description of the expected outputs and cross-references to the tutorial illustrating and explaining what to do in non-ideal situations. To ensure the reproducibility and the transparency of the TK analyses, MOSAIC_*bioacc*_ allows downloading all outputs under different formats, as well as the entire R code used (Figure 3.G).

**Fig. 3.**
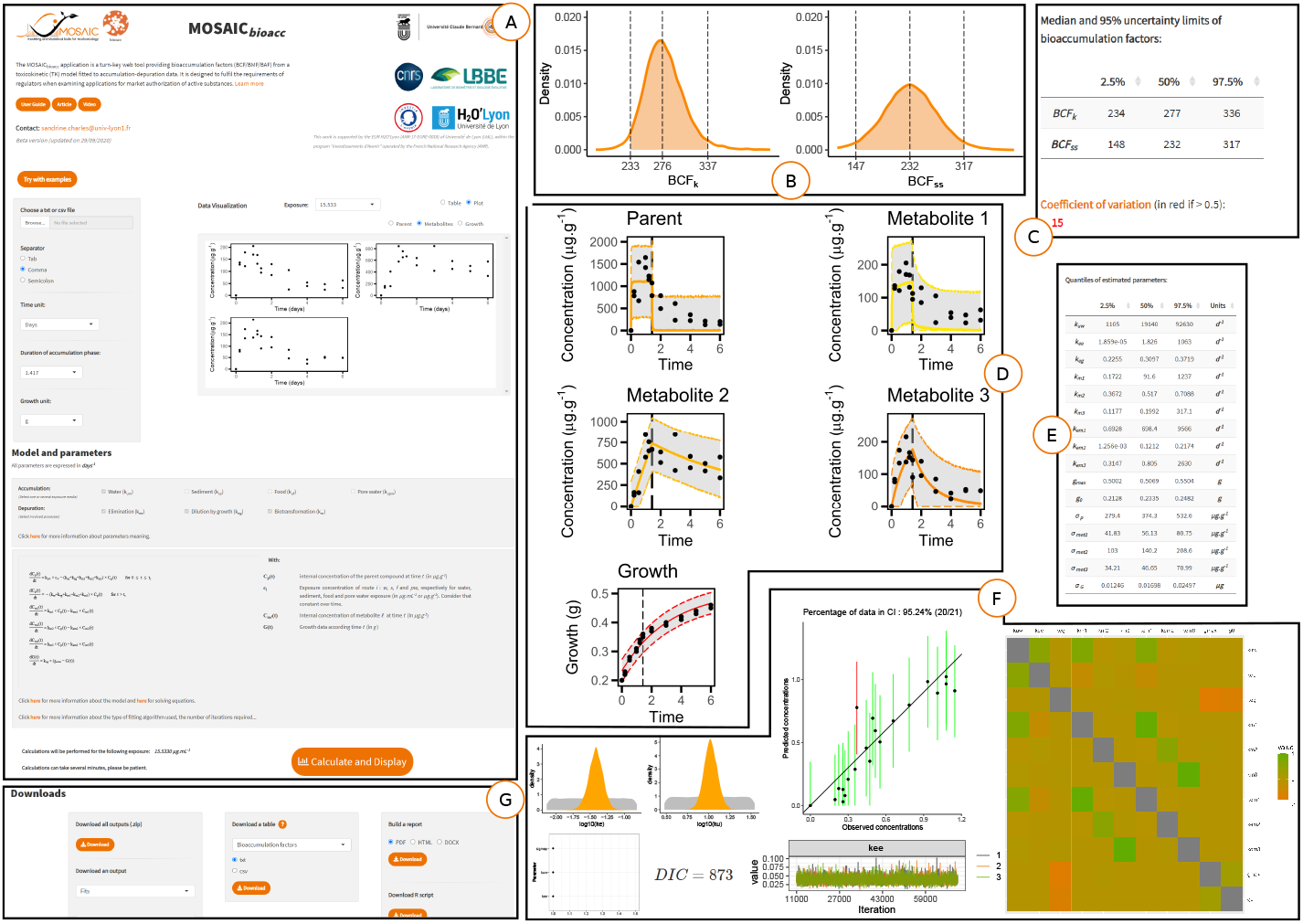
Selected pieces from MOSAIC_*bioacc*_ when performing a TK analysis on two sample data sets: `Oncorhynchus_two` and `Male_Gammarus_seanine`: (A) upload of experimental data and simplified summary of the TK model and its parameters (automatically delivered); (B) graphical representation of bioaccumulation factors (here the kinetic BCF with example `Oncorhynchus_two`); (C) the corresponding statistical summary of the BCF distribution; (D) TK model fit (concentration in the body as a function of time): median curve (solid colored line) and its uncertainty (gray area delimited by colored dotted lines); (E) estimation of model parameters fitted to bioaccumulation data; (F) various model goodness-of-fit criteria; (G) result downloading the results.

Several updates were recently implemented in MOSAIC_*bioacc*_. First, it is now possible to account for the lipid fraction within organisms in calcula-tions; users just need to enter their measured value. Secondly, MOSAIC_*bioacc*_ allows users to fit several nested TK models on a same data set. In practice, users just need to choose the parameters they want to appear in sub-models. According to the experimental conditions, several sub-models can indeed be considered and compared depending on the hypotheses to test either on the exposure routes or on the elimination processes. As illustrated in a case study in supplementary information (see full report in SI), organisms may have been exposed via several media (water and sediment in the case study in SI). By default, MOSAIC_*bioacc*_ fits the full TK model. Then users can test different TK sub-models, for example sub-models with only one exposure route (water or sediment in the case study in SI), and compare them to the full model based on both the Deviance Information Criteria (DIC) and the Watanabe–Akaike information criterion (WAIC) delivered by MOSAIC_*bioacc*_. Users can also test different TK sub-models ignoring some of the elimination processes even if they have been measured (*e*.*g*., neglecting the dilution by growth). Hence, users have now the possibility to choose the most appropriate TK model regarding their data. Third, a collection of more than 80 data sets is made available to support all features of MOSAIC_*bioacc*_. More than 95% of these data sets are published in the scientific literature. They encompass more than 25 species (aquatic, terrestrial, insect), more than 66 chemical substances, different exposure routes (water, sediment, soil, food) and several elimination processes (biotransformation and growth dilution). This data collection is presented as a table that summarises the main characteristics of the data (genus, category, substance, accumulation duration, exposure routes, number of data and replicates), as well as a direct link to the reference, and direct links to download the raw data and the full report provided by MOSAIC_*bioacc*_. In addition, the table also gives the kinetic bioaccumulation factor estimate (as a median and a 95% uncertainty interval).

### 3.2 GUTS models

#### 3.2.1 Few words about GUTS modelling

All GUTS models are today unified within a theoretical framework describing stressor effects on survival over time, based on hypotheses related to the stressor quantification, the compensatory processes (such as recovery), and the nature of the death process (Jager & Ashauer, 2018). In support of ERA, EFSA considers that both reduced versions of GUTS models (GUTS-RED models) are ready-to-use (Ockleford et al., 2018). To write it simple, these two reduced versions can only be used with standard toxicity test data, that is without measurements of internal damages within organisms. The SD version (the GUTS-RED-SD model) assumes that all individuals are identically sensitive to the chemical substance by sharing a common internal threshold concentration and that death is a stochastic process once this threshold is exceeded. The GUTS-RED-SD model then describes the instantaneous hazard rate as a threshold function of the damages, themselves described by a very simple TK model. The IT version (the GUTS-RED-IT model) is using the same TK part as the GUTS-RED-SD version. For its TD part, it is based on the critical body residue approach, which assumes that individuals differ in their tolerance threshold when exposed to a chemical compound according to a probability distribution. The GUTS-RED-IT model also assumes that individuals die as soon as their internal concentration reaches their individual-specific threshold. By default, the between-individual variability is described by a log-logistic probability distribution.

In its recent scientific opinion (Ockleford et al., 2018), EFSA clearly states its support for the use of TKTD models at Tier-2 of ERA according to a specific workflow. Applied in particular for GUTS-RED models, this workflow consists in the following three steps: (1) **Calibration**, which consists in fitting both GUTS-RED models to toxicity test data collected at constant concentration under a standard protocol, in order to get parameter estimates associated with their uncertainty; (2) **Validation**, which consists in simulating the number of survivors over time, using both GUTS-RED models and the previously estimated parameters, but for time-variable exposure profiles under which data have also been collected. The simulated numbers of survivors for both models are then compared to observed ones and the prediction-observation adequacy is checked according to one visual validation criterion together with three quantitative validation criteria. These validation criteria were defined by EFSA with the perspective to choose the most appropriate model for the next step; (3) **Prediction**, which consists in simulating the survival probability over time with the previously chosen model and the parameter estimates obtained in step (1), for environmentally realistic exposure scenarios in order to assess risk on how far is the exposure profile from causing a pre-defined effect. Namely, this third step aims at determining the *x*% Lethal Profile (denoted *LP*_*x*_), that is the multiplication factor leading to an additional *x*% of reduction in the final survival rate at the end of the exposure. The next subsection guides the reader step by step to perform the EFSA workflow directly using MOSAIC.

#### 3.2.2 MOSAIC_GUTS−fit_

MOSAIC offers two services related to the use of GUTS-RED models to analyze standard survival data as function of both time and exposure concentration: MOSAIC_*GUTS−fit*_ for step (1) and MOSAIC_*GUTS−predict*_ for steps (2) and (3). All features of MOSAIC_*GUTS−fit*_ have already been detailed in Baudrot et al. (2018b). We just recall here the main highlights: a facilitated uploading of data (either from example data files or from the users themselves), an automatic GUTS fitting analysis for either GUTS-RED-SD and GUTS-RED-IT models, all useful fitting outputs to check the relevance of the results (parameter estimates, fitting curve with its uncertainty, posterior predictive check), and a collection of *LC*_*x*_ calculations associated with their uncertainty (Figure 4.A). In the following subsection, MOSAIC_*GUTS−predict*_ is presented in details, in support of the validation and the prediction steps of the EFSA workflow.

**Fig. 4.**
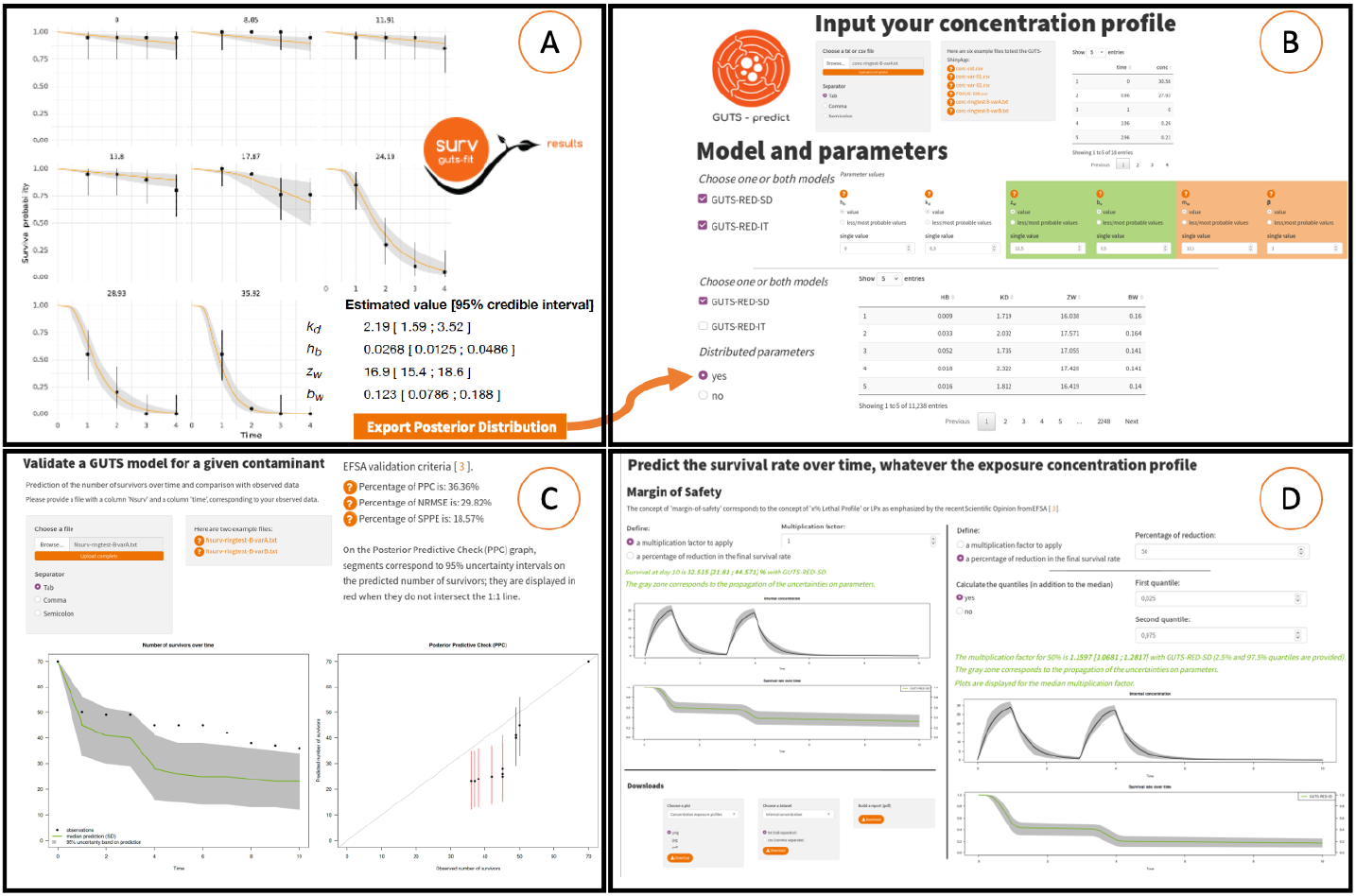
Selected pieces from MOSAIC for GUTS models: (A) GUTS **calibration** results: model predictions superimposed to the data, parameter estimates and the way to download the joint posterior distribution, from file `Ring-test Dataset B-cst`; (B) GUTS-predict first panel to enter the exposure profile for the simulation (EFSA steps (2) and (3), from file `conc-ringtest-B-varA.txt`), as well as to choose the model to use and how to consider its parameters (distributed or not, from file `mcmc-ringtest-B-SD.txt`); (C) outputs of the EFSA **validation** step (2) where the predicted number of survivors is compared to observed data (from file `Nsurv-ringtest-B-varA.txt`), together with EFSA validation criteria values; (D) outputs of the EFSA **prediction** step (3) where two options are proposed to quantify how far is the exposure profile from causing an *x*% effect: fixing the multiplication factor and simulating the predicted survival over time, or fixing *x* and getting the corresponding multiplication factor; and (E) downloading panel of MOSAIC_*GUTS−predict*_.

#### 3.2.3 MOSAIC_GUTS−predict_

MOSAIC_*GUTS−predict*_ has been developed in R (R Core Team, 2021) within a Shiny environment (Chang et al., 2021). It is available at https://mosaic.univ-lyon1.fr/guts-predict and performed using the computing facilities of the CC LBBE/PRABI. Both steps (2) and (3) of the EFSA workflow require a time-variable exposure profile that needs to be uploaded first (Figure 4.B). Then the user can perform simulations with one or both GUTS-RED models, for which parameter values need to be entered (Figure 4.B). Regarding parameter values, two options are proposed: only point values (such as means, medians…) or distributed parameters, namely coming from MOSAIC_*GUTS−fit*_ as the joint posterior distribution, downloadable in advance (Figure 4.A). From here, users can perform validation step (2) to predict the number of survivors over time to be compared with observed data (‘Validation’ tab). For this step, MOSAIC_*GUTS−predict*_ expects to receive as input both distributed parameters (in order to propagate the uncertainty all along the simulation) and a data file with observations under the uploaded exposure profile (typically a pulsed exposure, Figure 4.C). MOSAIC_*GUTS−predict*_ returns EFSA validation criteria values together with the simulation superimposed to the observed data and the posterior predictive check (PPC) graph. In the following or independently, the prediction step (3) can be performed to predict the survival probability over time as a function of time under the previously (or a new one) uploaded exposure profile. Usually, for step (3), users are using realistic scenarios, for example predicted environmental concentrations of active substances of plant protection products (European Food Safety Authority, 2017). This prediction step (‘Prediction’ tab) also requires the use of distributed parameters (namely according to their joint posterior distribution, as delivered in step (1)). From here, users have two options: (i) to fix a multiplication factor (MF) to apply on the uploaded exposure profile and get the prediction as a curve (the median tendency and its uncertainty) associated with the predicted survival probability at final time; (ii) to fix a percentage of additional reduction on survival at final time (*e*.*g*., 20% as on Figure 4.D, left) and ask MOSAIC_*GUTS−predict*_ to return the corresponding MF that could be applied with a *x*% of risk in terms of survival probability for the species-compound combination under interest; this MF is exactly the newly concept of the *x*% Lethal Profile (*LP*_*x*_) as defined by EFSA (Ockleford et al., 2018). Finally, users can download selected pieces of results (Figure 4.D, right).

## 4 New perspectives for Tier-1 in ERA

As detailed above, TKTD models allow to account for both time and concentration in predicting effects due to chemical exposure. In essence, based on standard protocols, TKTD models benefit from all collected data, while dose-response models only rely on data at a fixed target time, that is one of the time points in the experimental design, the most often the end of the experiment. Starting from the hypothesis that the gain in knowledge in using TKTD models allow a better precision (or, equivalently, a reduced uncertainty) on parameter estimates, this section highlights the added-value of GUTS models for the estimation of lethal concentration as required for Tier-1 in ERA.

As detailed in Baudrot & Charles (2019), the lethal concentration can be obtained from a GUTS model as a continuous function of both the chosen percentage *x* and the exposure duration *t* according to model parameter estimates. Hence, the calculation of any *LC*_(*x,t*)_ can be associated to its uncertainty, by propagating the uncertainty associated to model parameter estimates as accessible from the joint posterior distribution after performing Bayesian inference. Based on a battery of 20 data sets, the classical *LC*_50_ value at final time, as estimated by a 3-parameters log-logistic model (equation (2)) is compared to the corresponding calculations obtained from both GUTS-RED-SD and GUTS-RED-IT models. The 20 data sets are standard survival data sets with a first set for 10 different species exposed to chlorpyriphos (Rubach et al., 2012), a second for species *Daphnia magna* exposed to seven veterinary antibiotics (Wollenberger et al., 2000) and three other data sets (Forfait-Dubuc et al., 2012). Each data set was fitted with the three models thanks to the R package morse (Baudrot et al., 2021). Note that the entire analysis can be identically reproduced using the MOSAIC platform. For each data set, goodness-of-fit criteria were good enough to support the relevance of the results (see example on Figure 5.A for the data set of *D. magna* exposed to potassium dichromate). So, for each data set, the *LC*_(*x,t*)_ estimates (as medians and 95% uncertainty intervals) were collected for *x* = 50% and at the end of the experiment, directly from parameter estimates when using the 3-parameters log-logistic model (parameter *e* in equation (1) above), or asking for the calculation after predicting the dose-response curve with both GUTS-RED models (Figure 5.B).

**Fig. 5.**
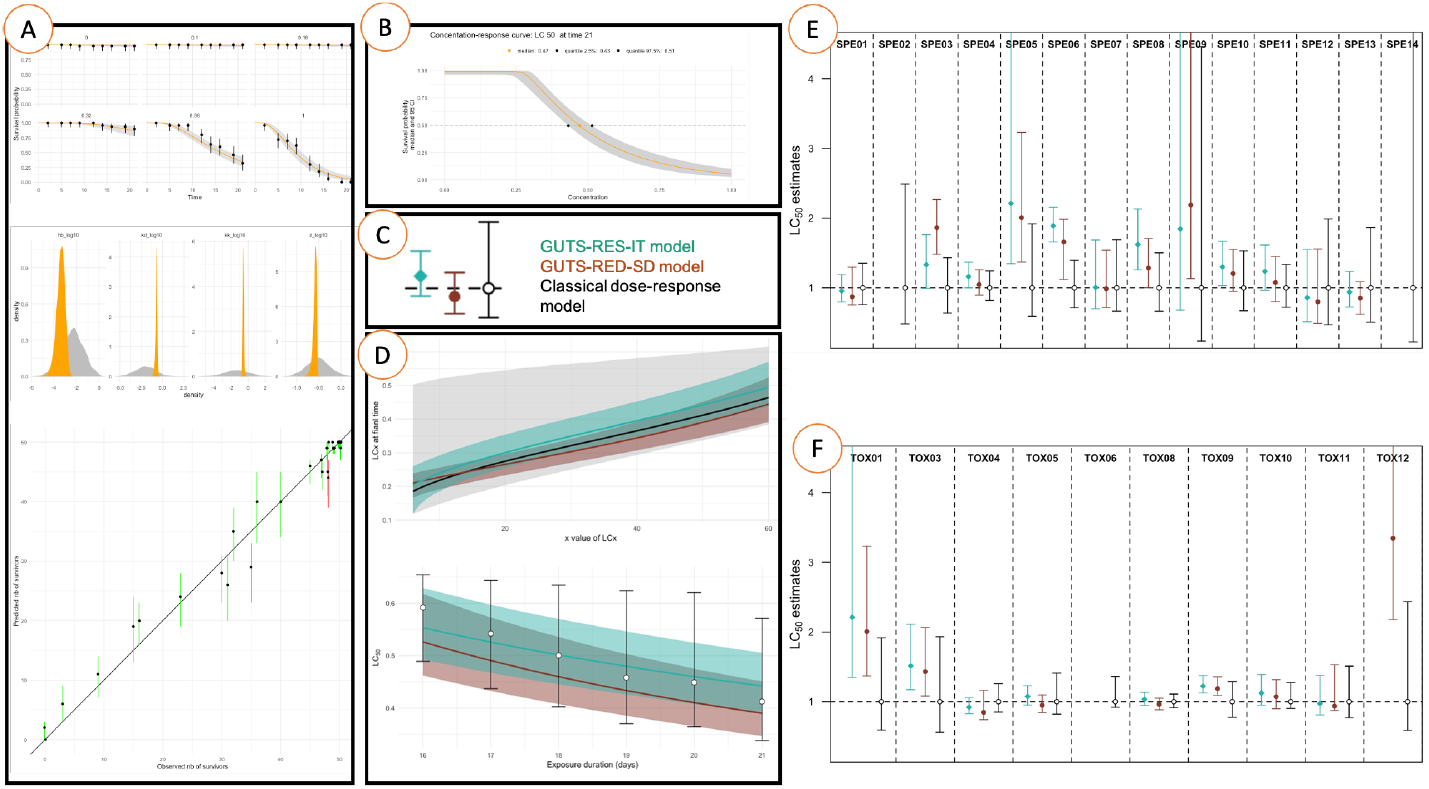
(A) GUTS-RED-SD fitting results for *Daphnia magna* exposed to potassium dichromate; (B) the corresponding predicted dose-response curve; (C) the three *LC*_50_ estimates with the 3-parameters log-logistic model (in black), the GUTS-RED-SD model (in red) and the GUTS-RED-IT model (in green); (D) *LC_x,t_* calculations for various *x* (upper panel) and various exposure time (lower panel); (E) comparison of *LC*_50_ estimates between species exposed to chlorpyriphos; (F) comparison of *LC*_50_ estimates between compound for *D. magna*.

Because of different orders of magnitude between *LC*_50_ estimates among data sets, the three *LC*_50_ estimates were compared by normalizing them to the classical *LC*_50_ median estimate obtained with the 3-parameters log-logistic model; this latter having thus a median of 1 (Figure 5.C). Focusing on the only data set of *D. magna* exposed to potassium dichromate (Figure 5.A-C), the starting hypothesis is confirmed with a better precision for both GUTS-RED estimates of the *LC*_50_, while both GUTS-RED estimates with a similar precision are not significantly different from the classical calculation (overlapping 95% uncertainty intervals). On the basis of this first finding, three questions deserve particular attention: (1) does the better precision depend on the *x* (first fixed at 50%)?; (2) does the better precision depend on the exposure duration (first fixed as the experiment duration)? (3) does the better precision depend on the data set, that is on the species? and/or on the compound? As shown on Figure 5.D for the combination *D. Magna*-potassium dichromate, the better precision does not depend neither on *x* nor on the time at which the *LC*_(*x,t*)_ is calculated. Figure 5.D also illustrates that *LC*_(*x,t*)_ estimates given by both GUTS-RED models can continuously be obtained whatever *t* between 0 and the exposure duration, but only at time points within the experimental design for the classical estimates with the 3-parameters log-logistic model.

Question (3) was answered in two steps. Figure 5.E first shows a slight dependency on the species exposed to chlorpyriphos of the *LC*_50_ precision, with a similar precision whatever the model for species 03 and 07, while both GUTS-RED estimates are different for species 03, 06 and 08. Secondly, Figure 5.F shows a slight dependency again, without high differences between both GUTS-RED estimates, but sometime different from the classic one (compound 03 and 09). These results need further investigation for example by looking at the phylogenetic proximity of the 10 compared species as well as at the mode of action of the seven compounds given that our knowledge is still poor in describing how effects vary across both species and compounds (Ashauer & Jager, 2018). Nevertheless, for most of the data set, both GUTS-RED models provide more precise *LC*_50_ estimates than a classical dose-response approach. Given that all facilities are today available to use GUTS models on standard data sets, the regulatory risk assessment should really consider the possibility to use them even at Tier-1.

## 5 Conclusions

Although tools are existing to use TKTD models, and although regulatory bodies strongly recommend their use for ERA (especially to facilitate the consideration of realistic exposure scenarios), practitioners struggle in appropriate them for reasons mostly attributable to modelers themselves. These reasons mainly come from lack of support: (1) to easily quantify uncertainties, and consequently their propagation to model outputs and subsequent predictions; (2) to better accept changing paradigm using new modelling approaches often appearing as black boxes, together with lack of support to fully perceive the concrete added-value of these novelties for their daily work; (3) to easily interpret goodness-of-fit criteria and therefore trust model results in their ability to support decisions from predictions; (4) to appropriate recent user-friendly turn-key facilities, while already recognized as automatically providing toxicity indices of interest in full compliance with regulatory guidelines and risk assessment decision criteria. The Bayesian inference framework is clearly the direction to take to facilitate the quantification of the uncertainties. In addition, practitioners will be most likely able to accept advanced modelling for ERA if accessibility of modelling is improved in terms of step-by-step support, reproducibility and transparency, founding principles of the web platform MOSAIC.

## 6 Ethical Approval

This article does not contain any studies with animals performed by any of the authors.

## 7 Consent to Participate

All authors participated in the research work underlying the content of the manuscript.

## 8 Consent to Publish

All authors read and approved the final manuscript for submission.

## 9 Authors Contributions

SC: coordinated part of the research work underlying the presented results as well as the writing of the manuscript with all contributors; she structured the final version of the manuscript, contributed to figures 2, 4 and 5, and conceived the graphical-art figure. AR: drafted the first version of manuscript, conceived figures 1 and 3, reviewed the manuscript and helped in finalizing the submitted version of the manuscript. VB: is the main developer of the morse package that supports the MOSAIC web interface; he is actively contributing to the MOSAIC_*GUTS−predict*_, and reviewed the manuscript. GM: developed the first version of MOSAIC_*bioacc*_ and made significant improvements in MOSAIC_*growth*_, then reviewed the manuscript. AS: is the main developer of MOSAIC_*GUTS−predict*_; she reviewed the manuscript and revised figure 4. DW: fully conceived MOSAIC_*growth*_ in its first version and revised the final manuscript. CL: coordinated part of the research work underlying the presented results and revised the entire manuscript.

## 12 Availability of data

All data used in this paper are downloadable from the MOSAIC web platform.

## Footnotes

1 To access to the very last version of MOSAIC_*bioacc*_ that is regularly updated and tested before to be deployed on the official server, please go to https://scharles-univlyon1.shinyapps.io/mosaic-bioacc-gamma/.

