## Supplementary material for "Taking full advantage of modelling to better assess environmental risk due to xenobiotics" (from another version of this preprint: version 2, 2021-03-31)

**Environmental Science and Pollution Research manuscript No.**
(will be inserted by the editor)

### Supplementary Information: Taking full advantage of modelling to better assess environmental risk due to xenobiotics

**Sandrine Charles**[*1] · **Aude Ratier**[*1] · **Virgile Baudrot**[1] · **Gauthier Multari**[1] · **Aurélie Siberchicot**[1] · **Dan Wu**[1] · **Christelle Lopes**[1]

Received: date / Accepted: date

*

These two authors equally contributed.

1

Université de Lyon, Université Lyon 1, CNRS UMR5558, Laboratoire de Biométrie et Biologie Evolutive, 69100 Villeurbanne, France.

### 1 Annex: Case study with sub-models

#### 1.1 Context

MOSAIC$_{bioacc}$ allows users to fit several nested TK models on a same data set. In practice, users just need to choose the parameters they want to appear in sub-models. According to the experimental conditions, several sub-models can indeed be considered and compared depending on the hypotheses to test either on the exposure routes or on the elimination processes. Organisms may have been exposed via several media (water and sediment in the following case study (section 1.2)). By default, MOSAIC$_{bioacc}$ fits the full TK model. Then users can test different TK sub-models, for example sub-models with only one exposure route (e.g., water or sediment (section 1.2)), and compare them to the full model based on both the Deviance Information Criteria (DIC) and the Watanabe–Akaike information criterion (WAIC) delivered by MOSAIC$_{bioacc}$.

#### 1.2 Case study data set

In order to illustrate this framework, the data set of *Physa acuta* exposed to AgNO3 spiked water and clean sediment for 7 days (Silva et al., 2020) was uploaded in MOSAIC$_{\mathrm{bioacc}}$.

##### *1.2.1 First hypothesis: complete TK model*

A first run of analyses was performed where parameters were automatically selected according to the corresponding experimental design (Eqs. S1 and S2), as given within the uploaded data set (i.e., water and sediment for the exposure routes).

$$\begin{aligned} \frac{dC_p(t)}{dt} &= k_{uw} \times c_w + k_{us} \times c_s - (k_{ee}) \times C_p(t) \quad \text{for } 0 \leq t \leq t_c \quad (S1) \\ \frac{dC_p(t)}{dt} &= -(k_{ez}) \times C_p(t) \quad \text{for } t > t_c \quad (S2) \end{aligned}$$

##### *1.2.2 Second hypothesis: only account for water exposure*

A second run of analyses was performed on the same data set that for the first run, but without considering the uptake rate from sediment (Eqs. S3 and S4), because the concentration of the chemical in the sediment is at environmental level.

$$\begin{aligned} \frac{dC_p(t)}{dt} &= k_{uw} \times c_w - (k_{ee}) \times C_p(t) \quad \text{for } 0 \leq t \leq t_c \quad (S3) \\ \frac{dC_p(t)}{dt} &= -(k_{ee}) \times C_p(t) \quad \text{for } t > t_c \quad (S4) \end{aligned}$$

#### 1.3 Results

For each analyses, the parameter estimates are summarized in Table 1. The WAIC and the DIC are penalized deviance statistics accounting for the number of parameters for use in model comparison for a same data set (Spiegelhalter et al., 2002). In this example (Table 1), the WAIC and DIC are similar between the two runs, as well as the estimations of $k_{ee}$, considering or not the sediment exposure route. Thus, for this data set, and applying the parsimony principle, it can be deduced that *P. acuta* accumulate AgNO3 principally by water. Silva et al. (2020) also concluded that when accounting for double exposure via both water and clean sediment, water was likely to be the main route.

***Table 1.*** *Summary of parameters estimated for the Physa_AgNO3_Silva2020.csv data set for run 1 (complete TK model) and for run 2 (without considering sediment exposure route, $k_{us}$ = 0). For parameters, the median value in given with its 95% credible interval into bracket.*

| | run 1 | run 2 |
|---|---|---|
| $k_{uw}$ (days$^{-1}$) | **1.492** [1.874*10$^{-5}$;60480] | **679**[568.9;972.6] |
| $k_{us}$ (days$^{-1}$) | **624.6**[30.6;903.6] | **0** |
| $k_{ee}$ (days$^{-1}$) | $\mathbf{5.605 * 10^{-3}}$ [1.421*10$^{-5}$;0.07212] | $\mathbf{3.952 * 10^{-3}}$ [1.382*10$^{-5}$;0.07106] |
| WAIC | 177.9 | 178 |
| DIC | 178.3 | 178.6 |

#### 1.4 Full reports

##### *1.4.1 Complete TK model (water, sediment and elimination)*

### MOSAIC$_{\text{bioacc}}$ REPORT

2021-03-02

---

This report is provided by the MOSAIC$_{\text{bioacc}}$ application available here:
https://mosaic.univ-lyon1.fr/bioacc

Contact:

MOSAIC$_{\text{bioacc}}$ uses the JAGS (version 4.3.0) and R (version 4.0.2) software, and in particular packages RJags (version 4.10), jagsUI (version 1.5.1) and Shiny (version 1.6.0).

The MOSAIC$_{\text{bioacc}}$ application is a turn-key web tool providing bioaccumulation factors (BCF/BSAF/BMF) from a toxicokinetic (TK) model fitted to accumulation-depuration data. It is designed to fulfil the requirements of regulators when examining applications for market authorization of active substances.

---

##### Data summary

File used: Physa_AGNO3_Silva2020.csv

Exposure: 0.0101 $\mu g.g^{-1}$

Accumulation phase duration: 7 days

Number of replicates: 3

Times: 0, 1, 2, 5, 7, 8, 9, 12, 14

Exposure routes: water Exposure routes: sediment

Elimination routes: excretion

##### Bayesian inference

Three MCMC chains were used to estimate model parameters.

Number of iterations: 176062

Thin: 47

### TK Model

The TK model used for these calculations was:

$$\frac{dC_p(t)}{dt} = k_{uw} \times c_w + k_{us} \times c_s - (k_{ee}) \times C_p(t) \quad for\ 0 \leq t \leq t_c$$

$$\frac{dC_p(t)}{dt} = -(k_{ee}) \times C_p(t) \quad for\ t > t_c$$

with:

$t$: time (expressed in days )

$t_c$: duration of the accumulation phase (expressed in days )

$C_p(t)$: internal concentration of the parent compound at time (expressed in $\mu g.g^{-1}$)

$k_{ee}$: elimination rates of excretion (expressed per days $^{-1}$)

$c_w$: exposure concentration of water route (expressed in $\mu g.mL^{-1}$)

$k_{uw}$: uptake rate of water exposure (expressed per days $^{-1}$)

$c_s$: exposure concentration of sediment route (expressed in $\mu g.g^{-1}$)

$k_{us}$: uptake rate of sediment exposure (expressed per days $^{-1}$)

#### Bioaccumulation factor calculation

##### Calculations

$$BCF_k = \frac{k_{uw}}{k_{ee}}$$

$$BCF_{ss} = \frac{C_p(t_c)}{c_w}$$

$$BSAF_k = \frac{k_{us}}{k_{ee}}$$

##### Bioconcentration factor (BCF)

##### $\mathbf{BCF_k}$ plot

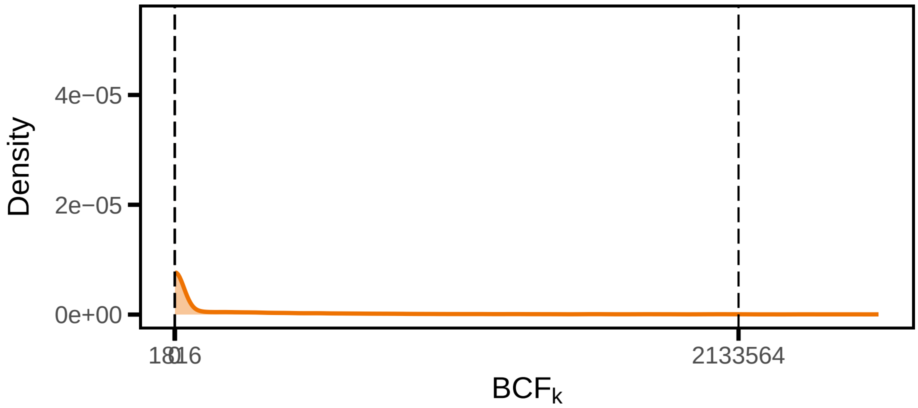


**$BCF_{ss}$ plot**

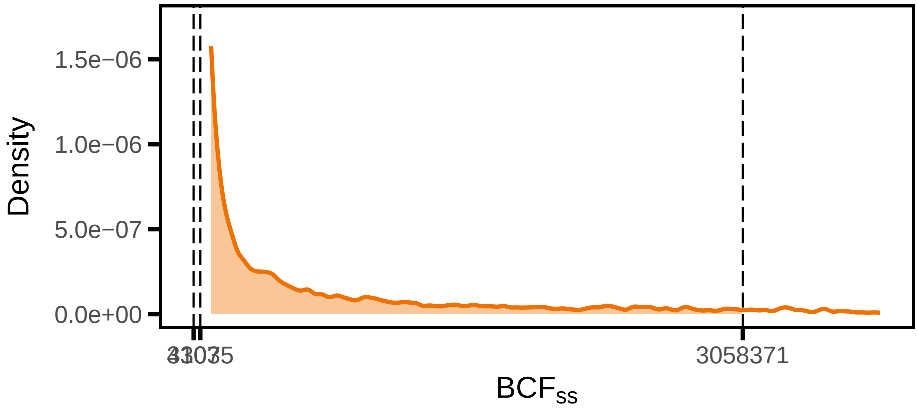


**BCF summary**

| | 2.5% | 50% | 97.5% | CV |
|---|---|---|---|---|
| BCFk | 0 | 1816 | 2133564 | 290 |
| BCFss | 3303 | 41075 | 3058371 | 19 |

**Biote-sediment accumulation factor (BSAF)**

**$BSAF_k$ plot**

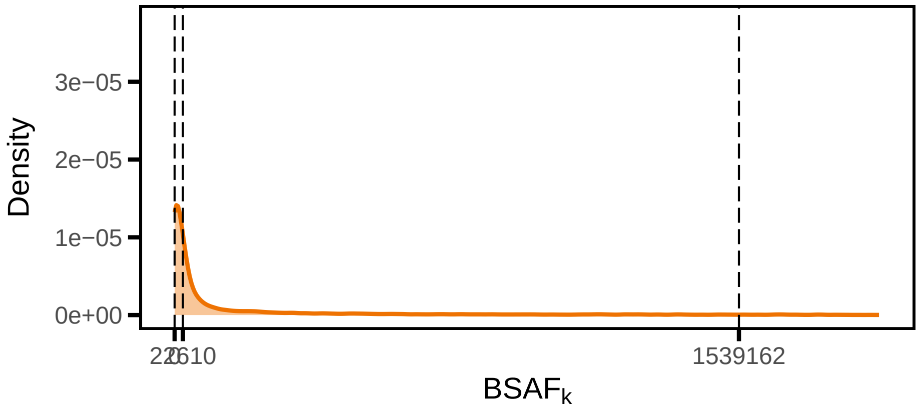


You didn't ask for the $BSAF_{ss}$

**BSAF summary**

| | 2.5% | 50% | 97.5% | CV |
|---|---|---|---|---|
| BSAFk | 0 | 22610 | 1539162 | 17 |

#### Fitting results

##### Fit plot

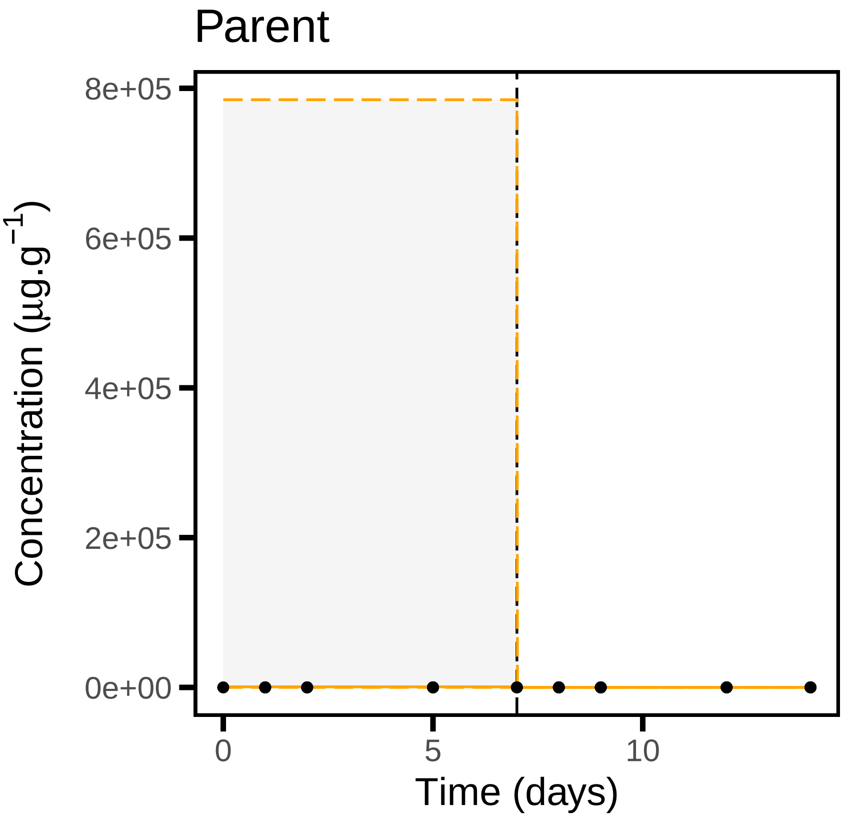

##### Quantiles of estimated parameters

| | 2.5% | 50% | 97.5% | |
|---|---|---|---|---|
| $k_{uw}$ | 1.874e-05 | 1.492 | 60480 | $d^{-1}$ |
| $k_{us}$ | 30.6 | 624.6 | 903.6 | $d^{-1}$ |
| $k_{ee}$ | 1.421e-05 | 5.605e-03 | 0.07212 | $d^{-1}$ |
| $\sigma_p$ | 8.434 | 11.18 | 15.84 | $\mu g.g^{-1}$ |

#### Goodness-of-fit criteria

##### Posterior Predictive Check

The PPC shows the observed values against their corresponding estimated predictions (black dots), along with their 95% credible interval (vertical segments). If the fit is correct, we expect to see 95% of the data within the intervals. Ideally observations and predictions should coincide, so we would expect to see black dots along the first bisector y = x (plain black line). The 95% credible intervals are colored in green if they overlap this line, in red otherwise.

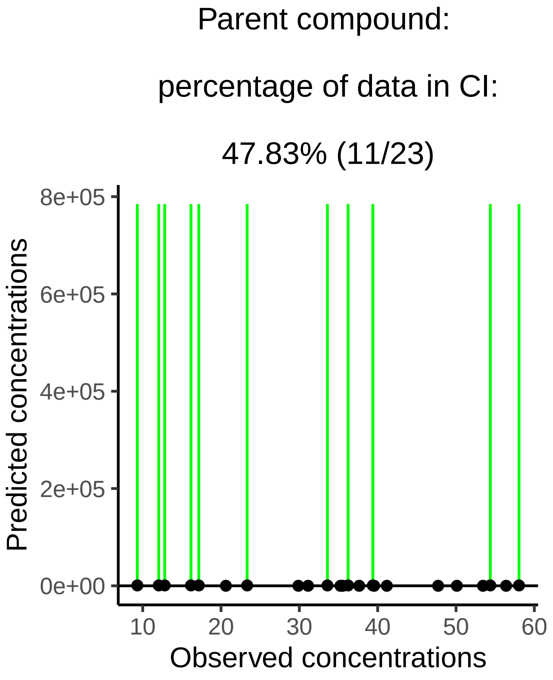


##### Priors and posteriors

The prior distribution is represented by the gray area and the posterior distribution by the orange area. The accuracy of the model parameter estimation can be visualized by comparing prior and posterior distributions: the overall expectation is to get a narrower posterior distribution compared to the prior one, what reflects that data contributed enough to precisely estimate parameters.

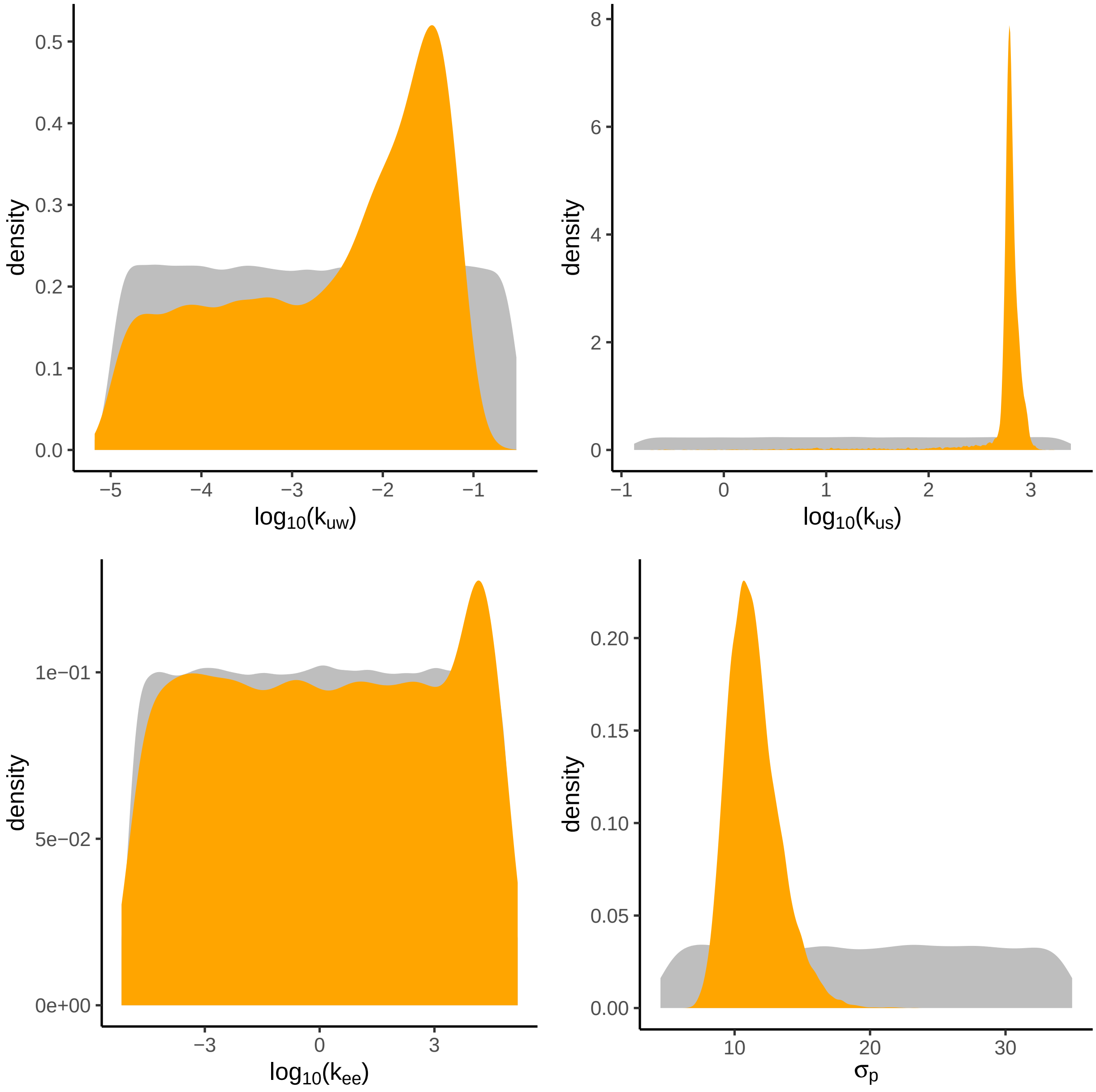


**Correlation between parameters**

If you want to see the coloured matrix giving a summary of parameter correlations, you need to import the corresponding figure directly from the application, page bottom, section "Downloads", then choose Download an output and select "GOF" then "parameter correlation". You can select the output format you prefer.

Correlations between parameters are visualized by projecting the joint posterior distribution in a plot matrix with planes of parameter pairs (lower triangular elements), marginal posterior distribution of each model parameter (diagonal), and Pearson correlation coefficients (upper triangular elements). Correlations are expected to be low (reflected by "potatoid" shapes of density lines in orange); a leaning elliptical shape translates high correlations (positive if leaning to the right, negative if leaning to the left).

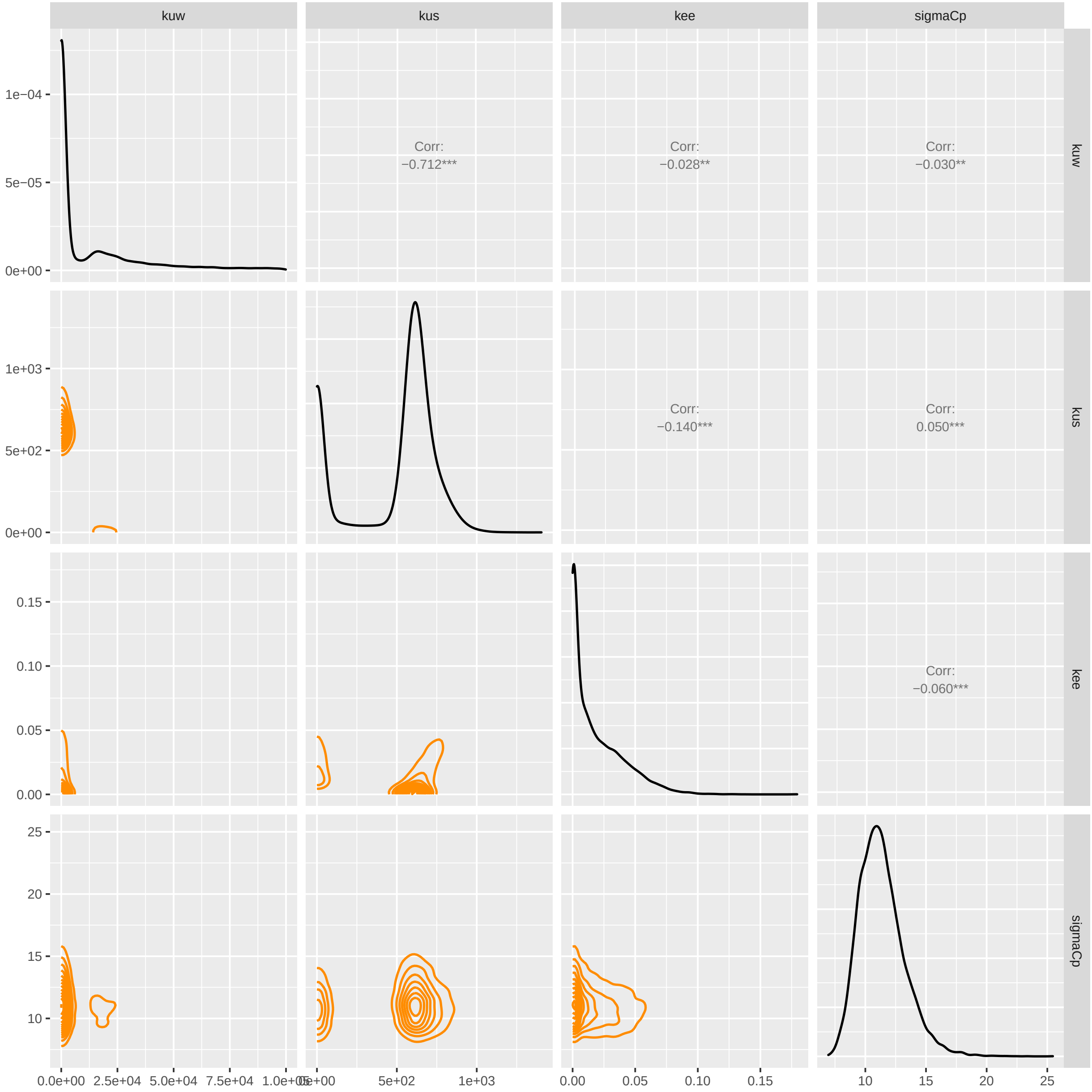


##### Potential Scale Reduction Factors

Convergence of the MCMC chains can be check with the Gelman-Rubin diagnostic expressed with the potential scale reduction factor (PSRF). Approximate convergence is diagnosed when the PSRF is below 1.01.

| | PSRF |
|---|---|
| $k_{uw}$ | 1.001 |
| $k_{us}$ | 1.002 |
| $k_{ee}$ | 1.000 |
| $\sigma_p$ | 0.999 |

##### Watanabe–Akaike information criterion

Information criteria offer a computationally appealing way of estimating the generalization performance of the model. A fully Bayesian criterion is the widely applicable information criterion (WAIC) by Watanabe a penalized deviance statistics accounting for the uncertainty in the parameters and can be used also for singular models. WAIC is widely used in model comparison for a same dataset (e.g., with or without $k_{ee}$). Sub-models with lower WAIC values will be preferred.

WAIC = 177.9

##### Deviance Information Criterion

This criteria, denoted DIC, is a penalized deviance statistics accounting for the number of parameters for use in model comparison for a same dataset (e.g., with or without $k_{ee}$). Sub-models with lower DIC values will be preferred.

DIC = 178.3

##### Traces of MCMC iterations

A traceplot is an essential plot for assessing convergence and diagnosing of MCMC chains. It shows the time series of the sampling process leading to the posterior distribution. Different colors are used for each of the chains (here 3) to assess within-chain convergence.

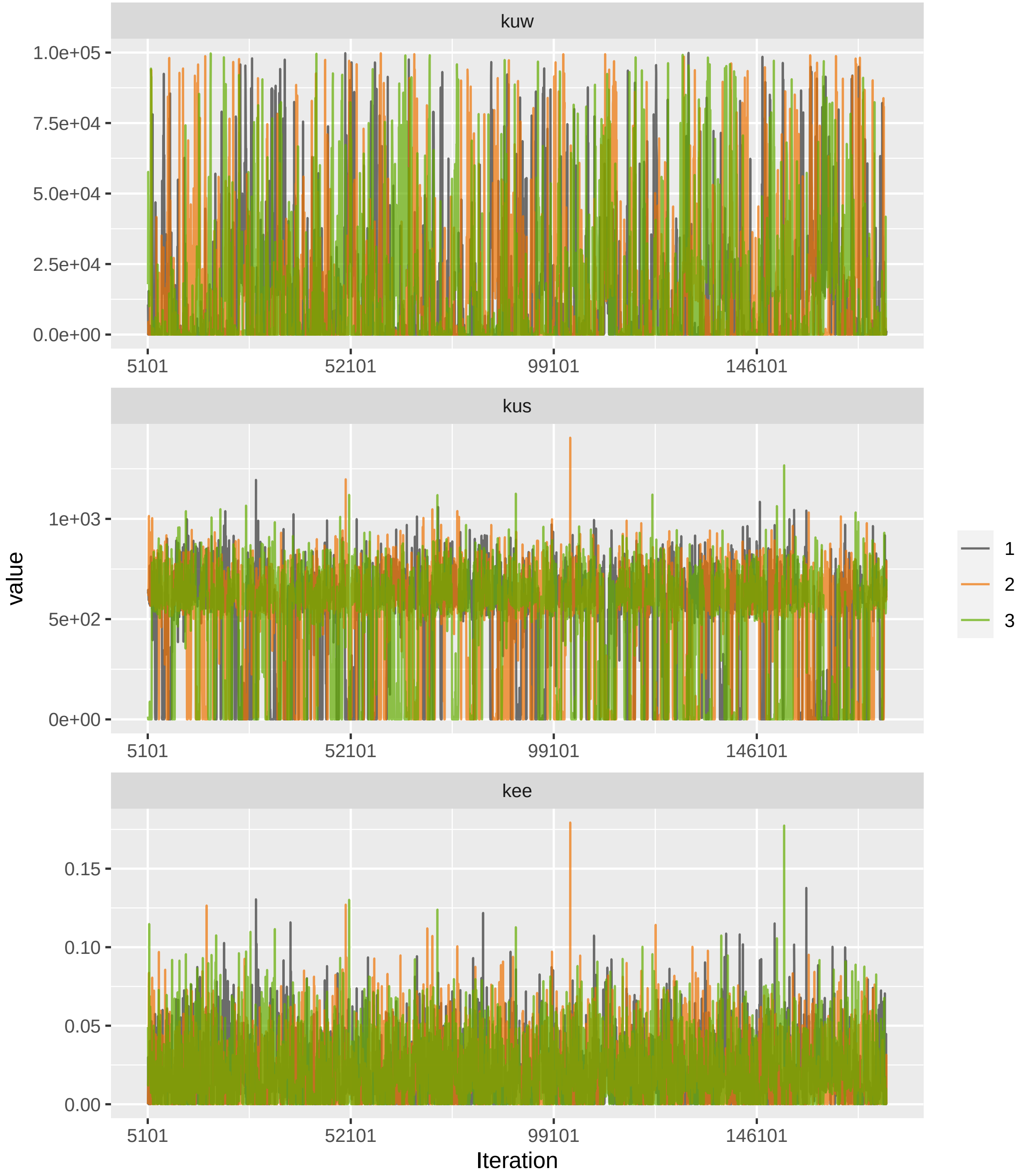
kuw
1.0e+05
7.5e+04
5.0e+04
2.5e+04
0.0e+00
5101
52101
99101
146101
kus
1e+03
5e+02
0e+00
kee
0.15
0.10
0.05
0.00
value
Iteration
1
2
3

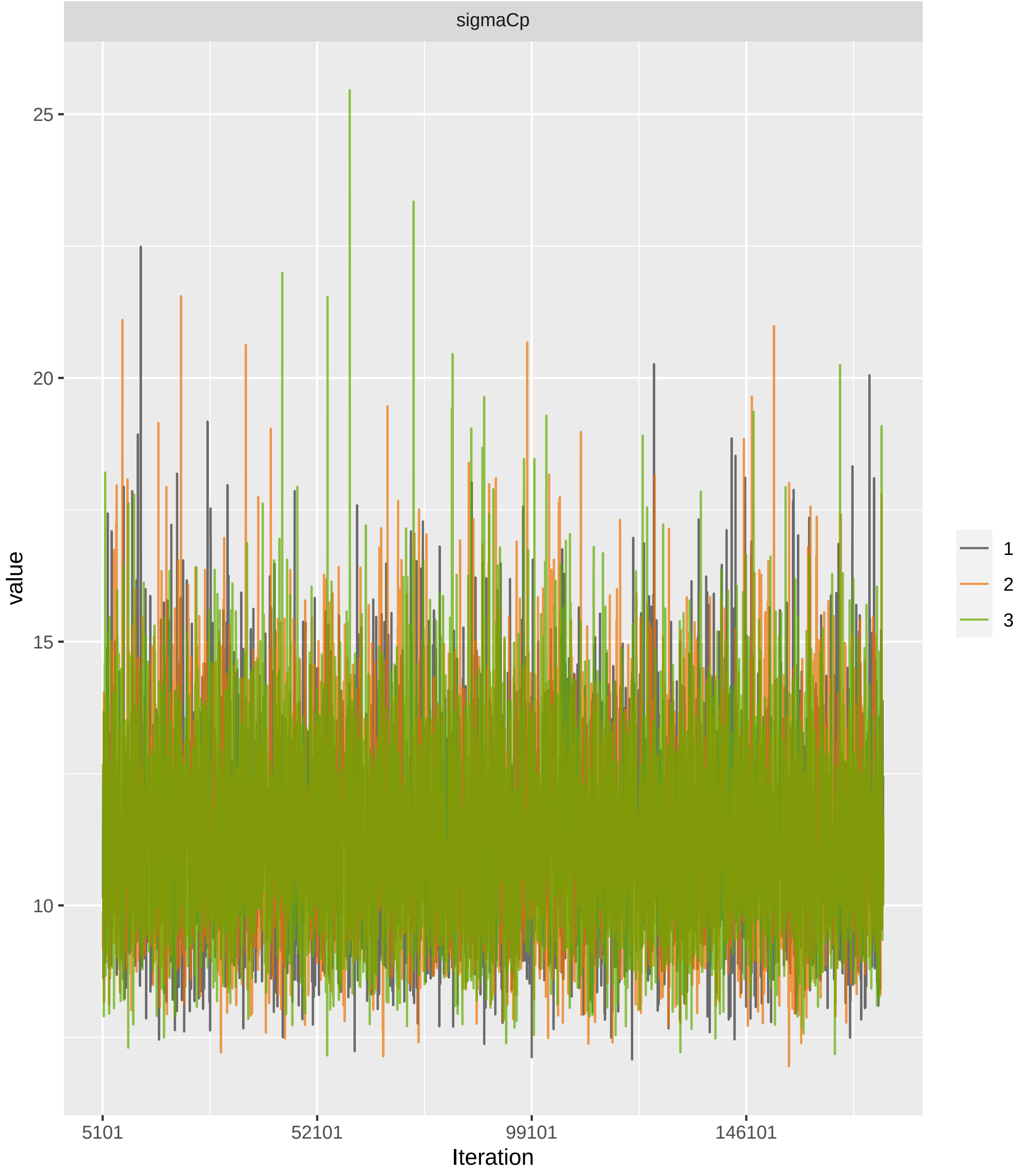
sigmaCp
25
20
15
10
value
5101
52101
99101
146101
Iteration
1
2
3

### Data Table

| time | conc | expw | exps | replicate |
|---|---|---|---|---|
| 0 | 0.435 | 0.0095 | 0.0101 | 1 |
| 1 | 12.811 | 0.0095 | 0.0101 | 1 |
| 1 | 12.050 | 0.0095 | 0.0101 | 2 |
| 1 | 17.158 | 0.0095 | 0.0101 | 3 |
| 2 | 23.338 | 0.0095 | 0.0101 | 1 |
| 2 | 16.165 | 0.0095 | 0.0101 | 2 |
| 2 | 9.317 | 0.0095 | 0.0101 | 3 |
| 5 | 36.227 | 0.0095 | 0.0101 | 1 |
| 5 | 39.379 | 0.0095 | 0.0101 | 2 |
| 5 | 54.379 | 0.0095 | 0.0101 | 3 |
| 7 | 58.044 | 0.0095 | 0.0101 | 2 |
| 7 | 33.587 | 0.0095 | 0.0101 | 3 |
| 8 | 56.398 | 0.0095 | 0.0101 | 1 |
| 8 | 39.550 | 0.0095 | 0.0101 | 2 |
| 8 | 29.876 | 0.0095 | 0.0101 | 3 |
| 9 | 53.448 | 0.0095 | 0.0101 | 1 |
| 9 | 41.165 | 0.0095 | 0.0101 | 2 |
| 9 | 20.621 | 0.0095 | 0.0101 | 3 |
| 12 | 31.118 | 0.0095 | 0.0101 | 1 |
| 12 | 37.640 | 0.0095 | 0.0101 | 2 |
| 12 | 35.249 | 0.0095 | 0.0101 | 3 |
| 14 | 50.109 | 0.0095 | 0.0101 | 1 |
| 14 | 47.718 | 0.0095 | 0.0101 | 2 |
| 14 | 35.543 | 0.0095 | 0.0101 | 3 |

##### *1.4.2 Only water exposure route (water and elimination)*

### MOSAIC$_{\text{bioacc}}$ REPORT

2021-03-02

---

This report is provided by the MOSAIC$_{\text{bioacc}}$ application available here:
https://mosaic.univ-lyon1.fr/bioacc

Contact:

MOSAIC$_{\text{bioacc}}$ uses the JAGS (version 4.3.0) and R (version 4.0.2) software, and in particular packages RJags (version 4.10), jagsUI (version 1.5.1) and Shiny (version 1.6.0).

The MOSAIC$_{\text{bioacc}}$ application is a turn-key web tool providing bioaccumulation factors (BCF/BSAF/BMF) from a toxicokinetic (TK) model fitted to accumulation-depuration data. It is designed to fulfil the requirements of regulators when examining applications for market authorization of active substances.

---

#### Data summary

File used: Physa_AGNO3_Silva2020.csv

Exposure: 0.0101 $\mu g.g^{-1}$

Accumulation phase duration: 7 days

Number of replicates: 3

Times: 0, 1, 2, 5, 7, 8, 9, 12, 14

Exposure routes: water

Elimination routes: excretion

#### Bayesian inference

Three MCMC chains were used to estimate model parameters.

Number of iterations: 14984

Thin: 4

### TK Model

The TK model used for these calculations was:

$$\frac{dC_p(t)}{dt} = k_{uw} \times c_w - (k_{ee}) \times C_p(t) \quad for\ 0 \leq t \leq t_c$$

$$\frac{dC_p(t)}{dt} = -(k_{ee}) \times C_p(t) \quad for\ t > t_c$$

with:

$t$: time (expressed in days )

$t_c$: duration of the accumulation phase (expressed in days )

$C_p(t)$: internal concentration of the parent compound at time (expressed in $\mu g.g^{-1}$)

$k_{ee}$: elimination rates of excretion (expressed per days $^{-1}$)

$c_w$: exposure concentration of water route (expressed in $\mu g.mL^{-1}$)

$k_{uw}$: uptake rate of water exposure (expressed per days $^{-1}$)

### Bioaccumulation factor calculation

#### Calculations

$$BCF_k = \frac{k_{uw}}{k_{ee}}$$

$$BCF_{ss} = \frac{C_p(t_c)}{c_w}$$

#### Bioconcentration factor (BCF)

#### $\mathbf{BCF_k}$ plot

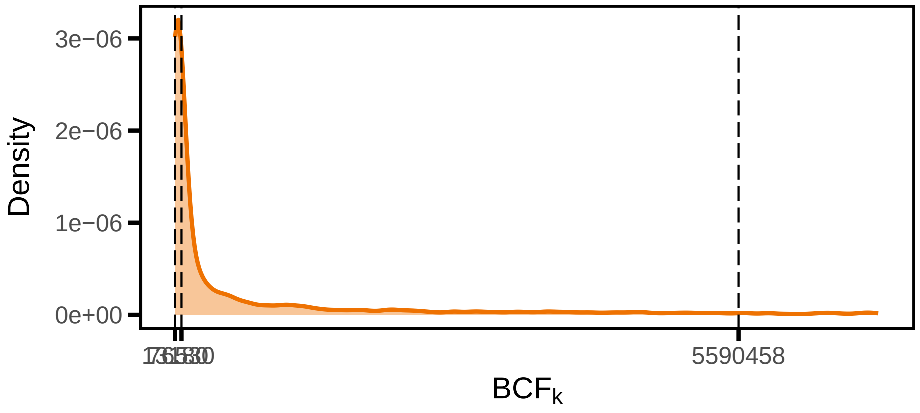

##### $BCF_{ss}$ plot

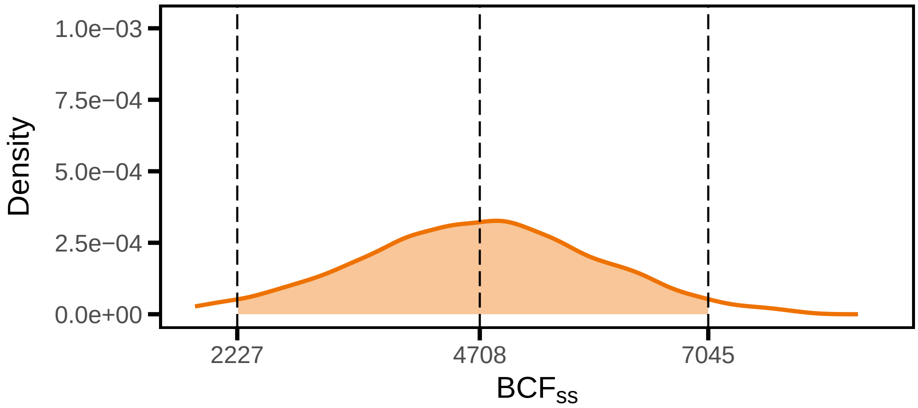


##### BCF summary

| | 2.5% | 50% | 97.5% | CV |
|---|---|---|---|---|
| BCFk | 13180 | 76530 | 5590458 | 18 |
| BCFss | 2227 | 4708 | 7045 | 0.26 |

#### Fitting results

##### Fit plot

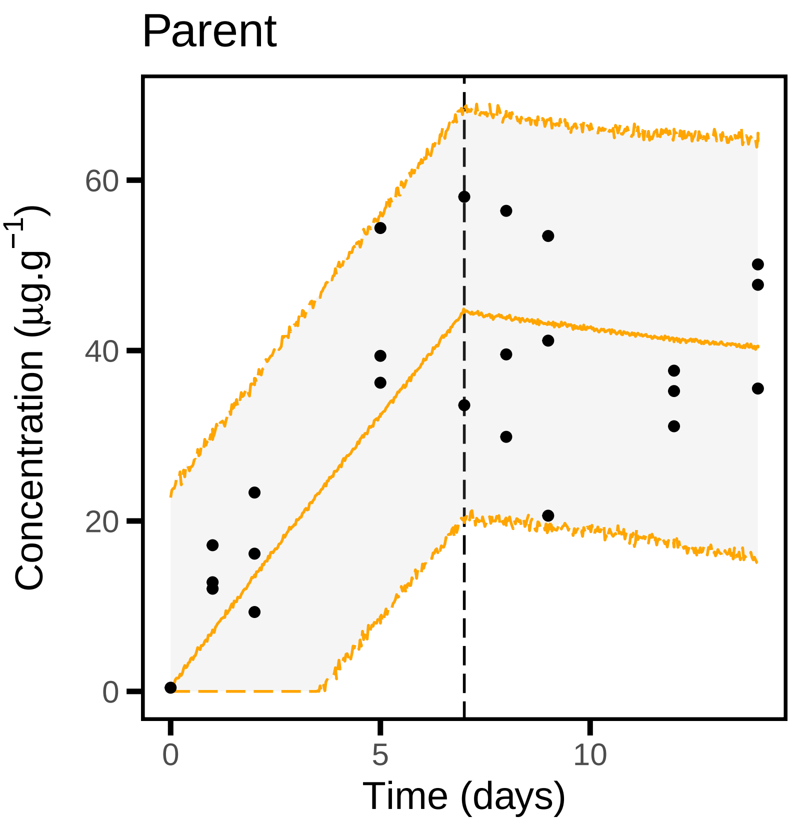


##### Quantiles of estimated parameters

| | 2.5% | 50% | 97.5% | |
|---|---|---|---|---|
| $k_{uw}$ | 568.9 | 679 | 972.6 | $d^{-1}$ |
| $k_{ee}$ | 1.382e-05 | 3.952e-03 | 0.07106 | $d^{-1}$ |
| $\sigma_p$ | 8.398 | 11.17 | 15.75 | $\mu g.g^{-1}$ |

#### Goodness-of-fit criteria

##### Posterior Predictive Check

The PPC shows the observed values against their corresponding estimated predictions (black dots), along with their 95% credible interval (vertical segments). If the fit is correct, we expect to see 95% of the data within the intervals. Ideally observations and predictions should coincide, so we would expect to see black dots along the first bisector y = x (plain black line). The 95% credible intervals are colored in green if they overlap this line, in red otherwise.

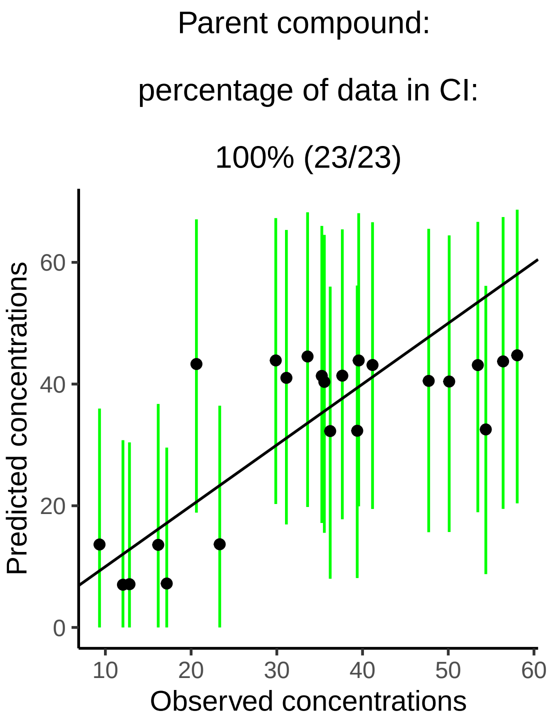


**Priors and posteriors**

The prior distribution is represented by the gray area and the posterior distribution by the orange area. The accuracy of the model parameter estimation can be visualized by comparing prior and posterior distributions: the overall expectation is to get a narrower posterior distribution compared to the prior one, what reflects that data contributed enough to precisely estimate parameters.

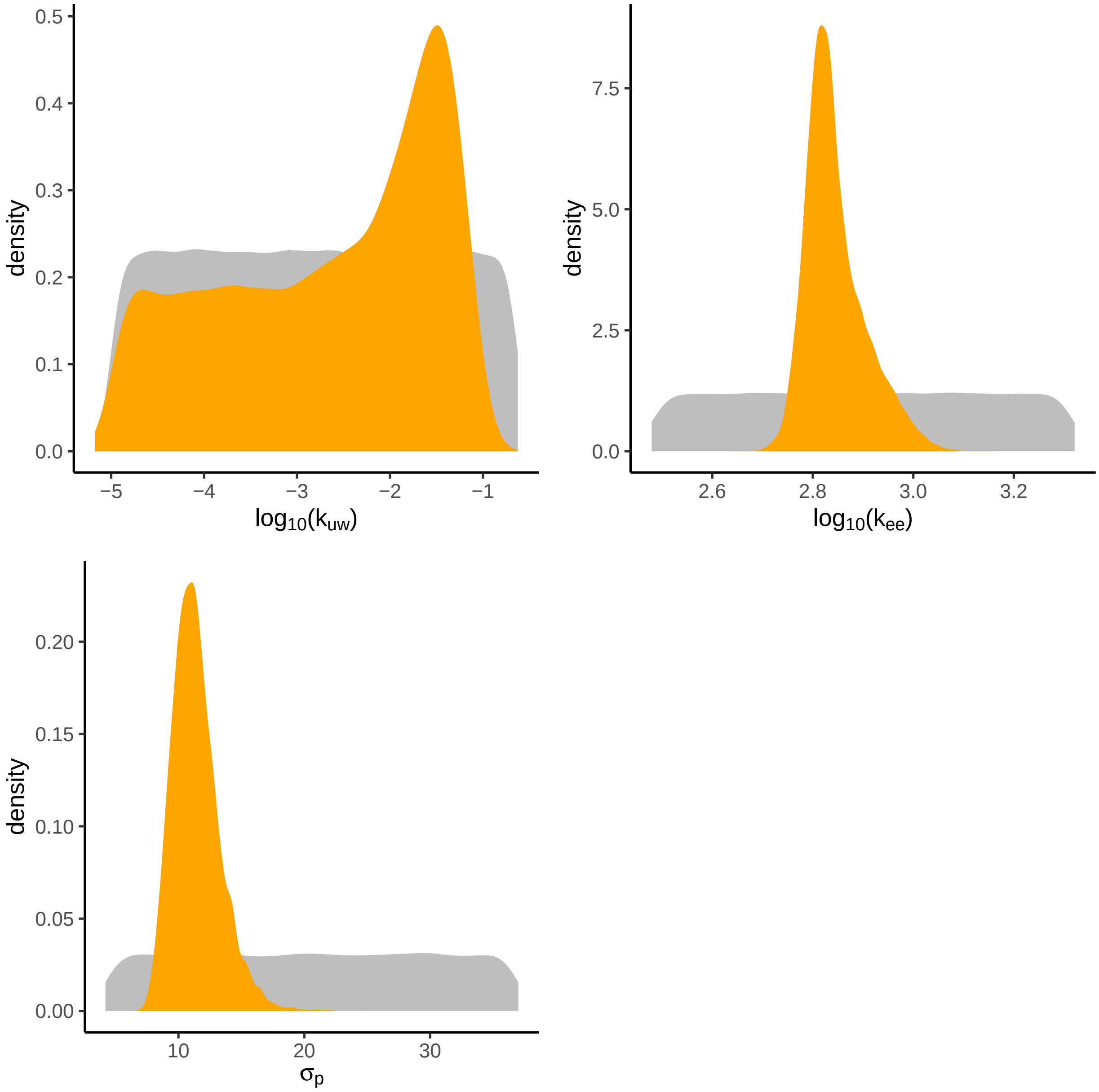


**Correlation between parameters**

If you want to see the coloured matrix giving a summary of parameter correlations, you need to import the corresponding figure directly from the application, page bottom, section “Downloads”, then choose Download an output and select “GOF” then “parameter correlation”. You can select the output format you prefer.

Correlations between parameters are visualized by projecting the joint posterior distribution in a plot matrix with planes of parameter pairs (lower triangular elements), marginal posterior distribution of each model parameter (diagonal), and Pearson correlation coefficients (upper triangular elements). Correlations are expected to be low (reflected by “potatoid” shapes of density lines in orange); a leaning elliptical shape translates high correlations (positive if leaning to the right, negative if leaning to the left).

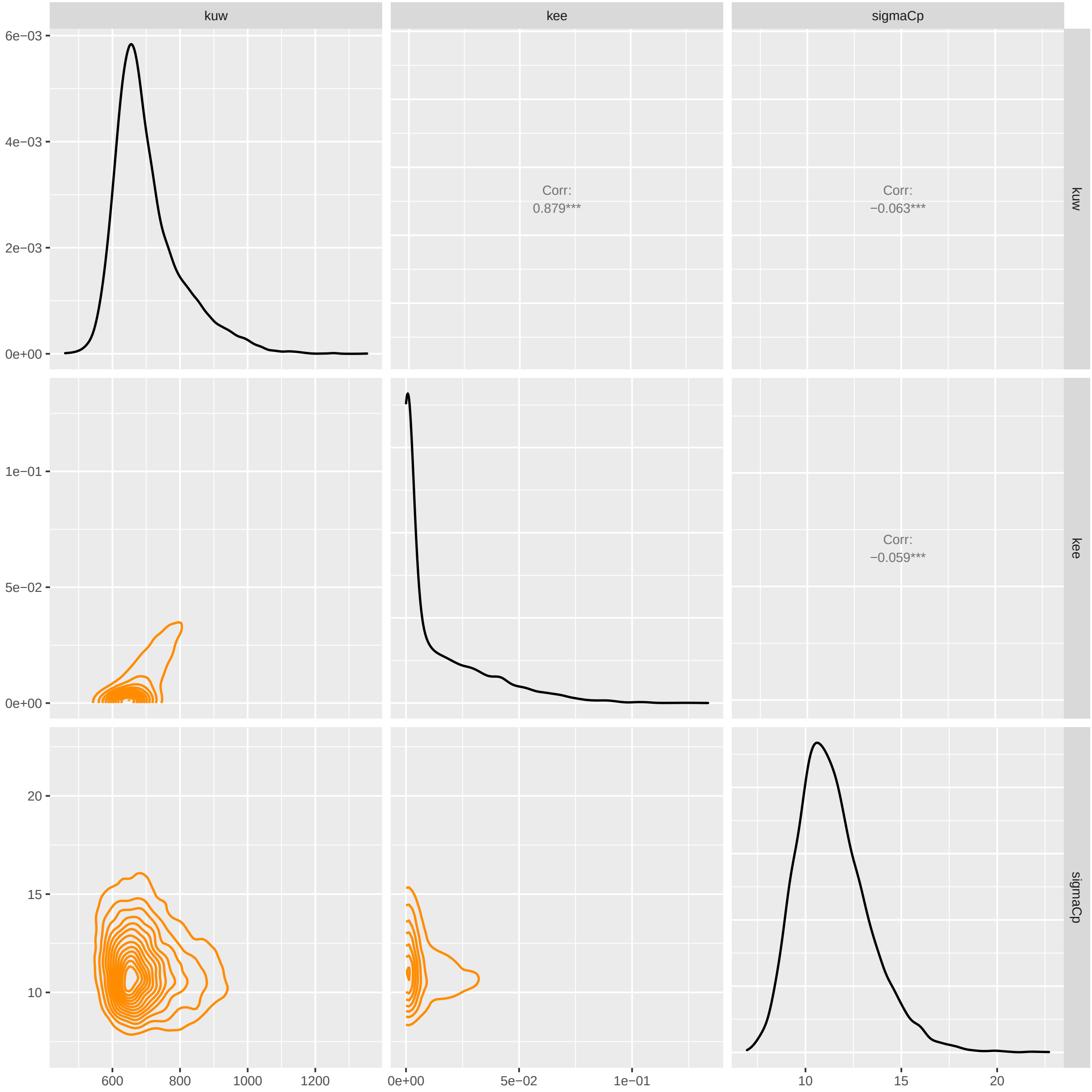


#### Potential Scale Reduction Factors

Convergence of the MCMC chains can be check with the Gelman-Rubin diagnostic expressed with the potential scale reduction factor (PSRF). Approximate convergence is diagnosed when the PSRF is below 1.01.

| | PSRF |
|---|---|
| $k_{uw}$ | 1.001 |
| $k_{ee}$ | 1.001 |
| $\sigma_p$ | 1.000 |

**Watanabe–Akaike information criterion**

Information criteria offer a computationally appealing way of estimating the generalization performance of the model. A fully Bayesian criterion is the widely applicable information criterion (WAIC) by Watanabe a penalized deviance statistics accounting for the uncertainty in the parameters and can be used also for singular models. WAIC is widely used in model comparison for a same dataset (e.g., with or without $k_{ee}$). Sub-models with lower WAIC values will be preferred.

WAIC = 178

**Deviance Information Criterion**

This criteria, denoted DIC, is a penalized deviance statistics accounting for the number of parameters for use in model comparison for a same dataset (e.g., with or without $k_{ee}$). Sub-models with lower DIC values will be preferred.

DIC = 178.6

**Traces of MCMC iterations**

A traceplot is an essential plot for assessing convergence and diagnosing of MCMC chains. It shows the time series of the sampling process leading to the posterior distribution. Different colors are used for each of the chains (here 3) to assess within-chain convergence.

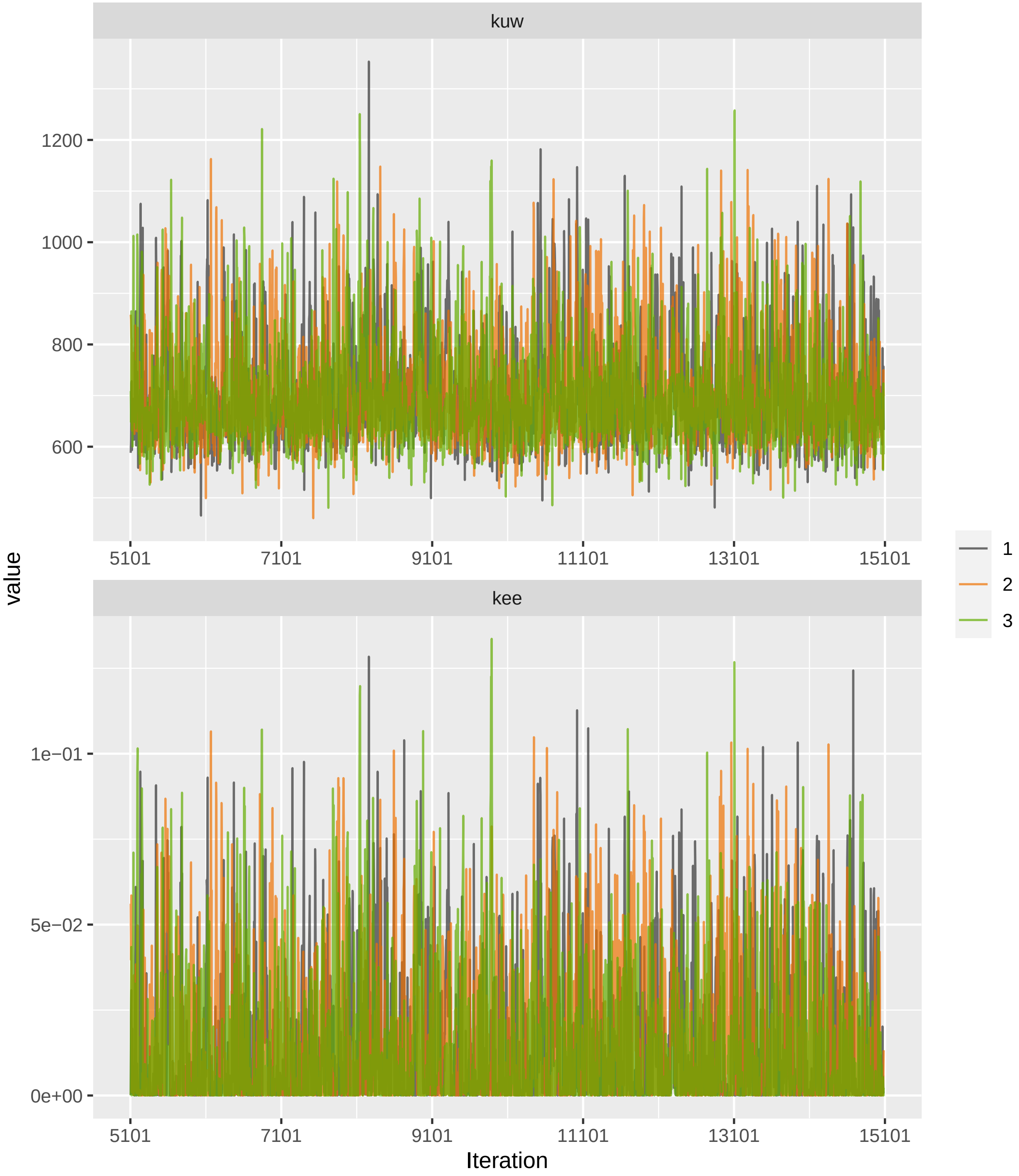
kuw
1200
1000
800
600
5101
7101
9101
11101
13101
15101
value
kee
1e−01
5e−02
0e+00
Iteration
1
2
3

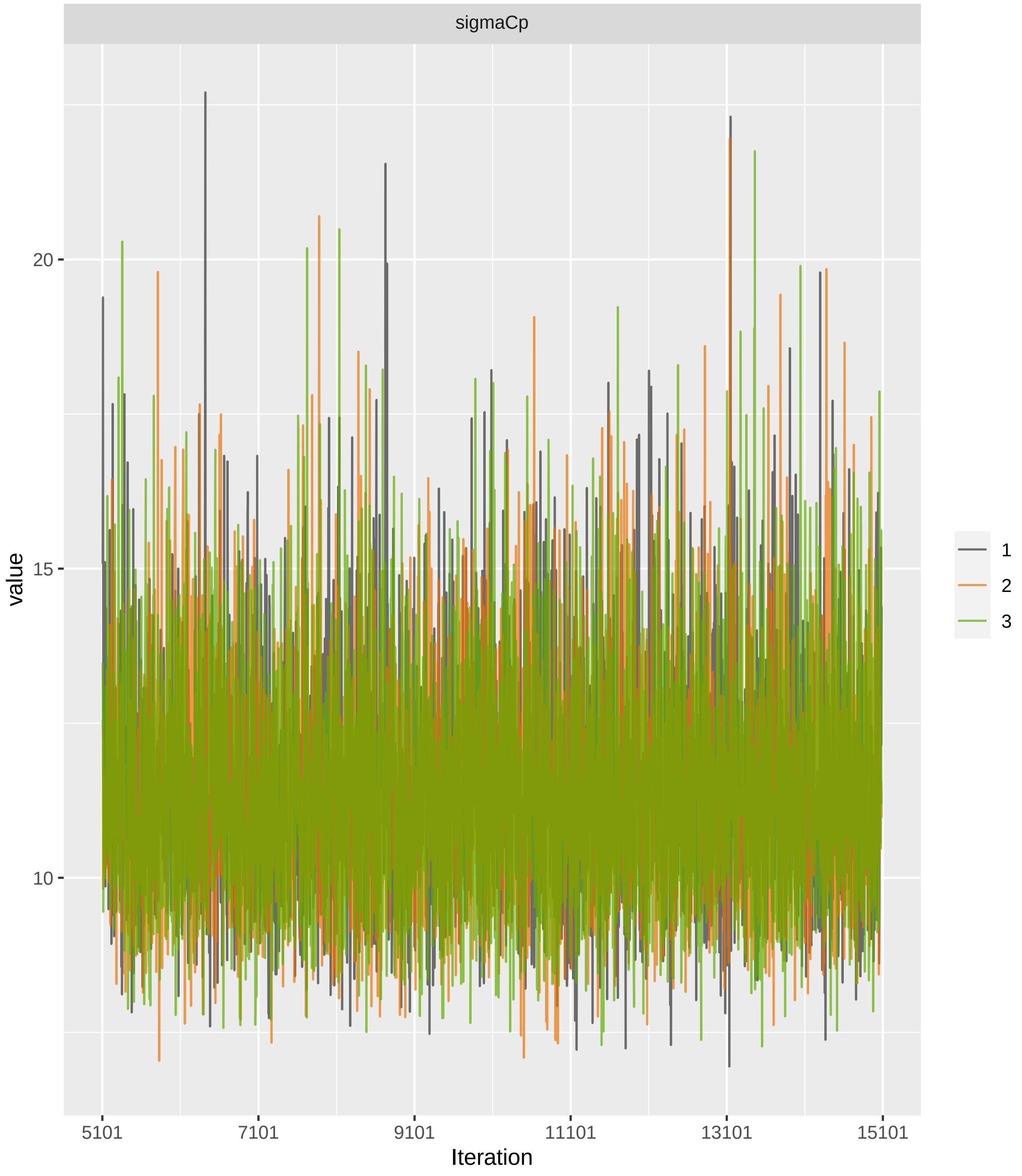
sigmaCp
value
20
15
10
5101
7101
9101
11101
13101
15101
Iteration
1
2
3

#### Data Table

| time | conc | expw | replicate |
|---|---|---|---|
| 0 | 0.435 | 0.0095 | 1 |
| 1 | 12.811 | 0.0095 | 1 |
| 1 | 12.050 | 0.0095 | 2 |
| 1 | 17.158 | 0.0095 | 3 |
| 2 | 23.338 | 0.0095 | 1 |
| 2 | 16.165 | 0.0095 | 2 |
| 2 | 9.317 | 0.0095 | 3 |
| 5 | 36.227 | 0.0095 | 1 |
| 5 | 39.379 | 0.0095 | 2 |
| 5 | 54.379 | 0.0095 | 3 |
| 7 | 58.044 | 0.0095 | 2 |
| 7 | 33.587 | 0.0095 | 3 |
| 8 | 56.398 | 0.0095 | 1 |
| 8 | 39.550 | 0.0095 | 2 |
| 8 | 29.876 | 0.0095 | 3 |
| 9 | 53.448 | 0.0095 | 1 |
| 9 | 41.165 | 0.0095 | 2 |
| 9 | 20.621 | 0.0095 | 3 |
| 12 | 31.118 | 0.0095 | 1 |
| 12 | 37.640 | 0.0095 | 2 |
| 12 | 35.249 | 0.0095 | 3 |
| 14 | 50.109 | 0.0095 | 1 |
| 14 | 47.718 | 0.0095 | 2 |
| 14 | 35.543 | 0.0095 | 3 |
